# Ramp sequence may explain synonymous variant association with Alzheimer’s disease in the Paired Immunoglobulin-like Type 2 Receptor Alpha (*PILRA*)

**DOI:** 10.1101/2025.01.06.631528

**Authors:** Justin B. Miller, J. Anthony Brandon, Lauren M. McKinnon, Hady W. Sabra, Chloe C. Lucido, Josue D. Gonzalez Murcia, Kayla A. Nations, Samuel H. Payne, Mark T.W. Ebbert, John S.K. Kauwe, Perry G. Ridge

## Abstract

**BACKGROUND:** Synonymous variant *NC_000007.14:g.100373690T>C* (*rs2405442:T>C*) in the Paired Immunoglobulin-like Type 2 Receptor Alpha (*PILRA*) gene was previously associated with decreased risk for Alzheimer’s disease (AD) in genome-wide association studies, but its biological impact is largely unknown.

**OBJECTIVE:** We hypothesized that *rs2405442:T>C* decreases mRNA and protein levels by destroying a ramp of slowly translated codons at the 5’ end of *PILRA*.

**METHODS:** We assessed *rs2405442:T>C* predicted effects on *PILRA* through quantitative polymerase chain reactions (qPCR) and enzyme-linked immunosorbent assays (ELISA) using Chinese hamster ovary (CHO) cells.

**RESULTS:** Both mRNA (*P*=1.9184 × 10^−13^) and protein (*P*=0.01296) levels significantly decreased in the mutant versus the wildtype in the direction that we predicted based on destroying a ramp sequence.

**CONCLUSIONS:** We show that *rs2405442:T>C* alone directly impacts *PILRA* mRNA and protein expression, and ramp sequences may play a role in regulating AD-associated genes without modifying the protein product.

**Research in Context:**

1. **Systematic review:** Genetic variants identified through genome-wide association studies often lack biological support for their association with Alzheimer’s disease. Although synonymous variant *rs2405442:T>C* in *PILRA* was previously reported as protective against Alzheimer’s disease, its effects have generally been attributed to linkage with missense variant, *rs1859788:A>G*.
2. **Interpretation:** We show that *rs2405442:T>C* alone decreases mRNA and protein levels by destroying a ramp of slowly translated codons at the beginning of *PILRA*. We also show that a ramp sequence is present in *PILRA* and likely regulates mRNA and protein levels, thus offering a plausible biological mechanism explaining *rs2405442:T>C* association with Alzheimer’s disease independent of *rs1859788:A>G*.
3. **Future directions:** We provide the first protocol to evaluate how disease-associated variants impact ramp sequences, which could explain why some genetic variants are reported by genome-wide association studies. Future studies might examine if the ramp sequence could be therapeutically targeted to regulate *PILRA* expression without changing the protein product.

---

Alzheimer’s disease (AD) is highly heritable, with genetic variants accounting for 58-79% of total dementia risk^1^. Common genetic effects identified through genome-wide association studies (GWAS) implicate approximately 80 genetic risk loci with AD-type dementia^2–9^, yet less is known about which genetic variants drive disease association. Many factors from high-impact diseases in addition to AD (i.e., amyloid plaques and neurofibrillary tangles) contribute to the dementia phenotype^10,11^, and heterogeneity plays a role in several distinct subtypes based on biomarkers^12–14^, genetics^15,16^, imaging^13,17–21^, and impact on daily function^22,23^. Similarly, clinical symptoms of dementia are heterogeneous and based on a progression of amyloid deposition, tau buildup, and neurodegeneration (A/T/N)^24^, with mixed pathologies impacting the speed of cognitive decline^13,17,22,25–29^. While polygenic risk scores (PRS) have recently emerged as a viable tool to aggregate genetic risk across various disease-associated loci so that complex population-specific genetic interactions can be simplified to a single risk score^30,31^, they do not attempt to characterize the biological mechanisms underpinning disease associations, and many associations have yet to be biologically validated. One of those currently unsubstantiated associations is located in the Paired Immunoglobulin-like Type 2 Receptor Alpha (*PILRA*). Here, we biologically assessed the effects of synonymous variant, *NC_000007.14:g.100373690T>C* (*rs2405442:T>C),* and propose that its association with AD is caused by destroying a ramp of slowly-translated codons at the 5’ end of *PILRA*.

Ramp sequences are essential genetic regulatory regions that counterintuitively maximize overall translational efficiency by slowing translation at the 5’ end of genes to evenly space ribosomes, which limits downstream ribosomal collisions and reduces translational errors^32–39^. Specifically, ramp sequences increase mRNA stability and gene expression, especially in genes that have higher ribosome density, higher mRNA levels, and a strong correlation between mRNA and protein expression^38,40^ by reducing ribosome stalling and mRNA degradation via ribosome-associated protein quality control (RQC)^41^. Ramp sequences are phylogenetically conserved^42^, yet differ between human populations^43^ and cell types^44^, which corresponds with population and cell-specific differences in gene expression^38,43,44^. A ramp sequence is present in *PILRA*, which likely helps regulate both protein and mRNA levels within different cell types.

*PILRA* is an inhibitory receptor that regulates immune cells^45^ of the myelomonocytic lineage such as macrophages, dendritic cells, monocytes and monocyte-derived dendritic cells and is highly expressed in the lymph node and neural tissues^46,47^. It functions by negatively regulating neutrophil infiltration and controlling monocyte mobility^45,48^. The innate and adaptive immune responses have been implicated in AD^49^, and gene regulation of *PILRA*-expressing myeloid cells have also associated with AD^50^. AD risk alleles are specifically enriched in active enhancers of myeloid-derived cells that express *PILRA* such as monocytes, macrophages, and microglia, with *PILRA* expression contributing a systemic failure of cell-mediated amyloid-β (Aβ) clearance^51^, which likely contributes to AD onset and progression.

Several studies have found AD-associated variants in *PILRA* to be protective^7,52,53^, yet the protective variant effects are generally attributed to a missense variant, *NC_000007.14:g.100374211A>G* (*rs1859788:A>G*)^53^, which is in high linkage disequilibrium with *rs2405442:T>C*. However, we show that *rs2405442:T>C* alone disrupts the *PILRA* ramp sequence by increasing codon adaptiveness relative to the rest of the transcript, which in turn significantly decreases both mRNA (*P*=1.9184 × 10^−13^) and protein (*P*=0.01296) levels in the direction that we hypothesized based on predicted ramp sequence effects. This study is the first time that ramp sequences have been used to prioritize disease-associated variants for biological validation and offers a likely biological mechanism that can regulate *PILRA* expression without altering the final protein product. Further, these analyses show that synonymous variant *rs2405442:T>C* alone disrupts *PILRA* and may drive association with AD.

## MATERIALS AND METHODS

### Identifying AD-associated genetic variants

We prioritized genetic variants for ramp sequence analyses using the GWAS summary statistics from Jansen, et al. ^54^ because they report all single nucleotide polymorphism (SNP) associations with AD that exceeded the genome-wide significance threshold of *P*≤5.0×10^−8^ before accounting for linkage disequilibrium at each locus. While Bellenguez, et al. ^2^ report additional genetic associations with AD, we opted to not include their summary statistics in these analyses because they report only variants that are likely independent hits after performing linkage disequilibrium analyses, which greatly reduces the number of reported genetic associations by using p-values to prioritize the independent hits. In some cases, leading variants are chosen based on predicted effects, which would also bias our analyses since variant-level ramp sequence effects have not previously been reported. Additionally, ramp sequences are affected by only exonic coding variants, which are generally rarer than intronic variants in GWAS and may be missed by a clumping approach to choose independent hits. Thus, we decided that the full table of variant-level associations reported by Jansen, et al. ^54^ was most appropriate to assess how ramp sequences potentially impact AD. Although 2,357 variants were originally reported^54^, only 51 variants were reported in exonic regions, with only 14 SNPs identified as plausible causal variants based on a fine-mapping model that accounts for *APOE* ε4, roughly corresponding to *P*>~2.0×10^−4^. All computational analyses were limited to those 14 variants.

Ensembl^55^ web queries were conducted in December 2023 to obtain the most severe variant consequence, highest minor allele frequency, Combined Annotation Dependent Depletion (CADD) score^56^, Genomic Evolutionary Rate Profiling (GERP) score^57^, and GRCh38.p14 reference genome chromosome coordinates. RegulomeDB^58^ scores were then queried using the RegulomeDB web interface. All transcript isoforms with GRCh38.p14 reference genome coordinates were downloaded from the National Center for Biotechnology Information (NCBI; https://www.ncbi.nlm.nih.gov/datasets/genome/GCF_000001405.40/) in December 2023.

### Identifying Ramp Sequences

Since ramp sequences are dependent on tissue and cell-specific tRNA pools, we used The Ramp Atlas^44^ to download pre-computed tRNA efficiency values for 62 human tissues included in a consensus dataset derived from the Genotype-Tissue Expression (GTEx)^59^, Functional ANnoTation Of the Mammalian genome (FANTOM5)^60^, and the Human Protein Atlas^47^ databases. An additional file consisting of codon efficiencies from 66 cell types was also downloaded from The Ramp Atlas and used to analyze cell-specific effects on *PILRA* ramp sequences. The relative codon adaptiveness in each tissue or cell type could impact the presence or absence of a ramp sequence by changing where translational bottlenecks occur without altering the DNA sequence. We used ExtRamp^36^ to identify ramp sequences in the reference and mutant sequence for each tissue or cell type individually using the `-a` option to specify the relative codon adaptiveness for each tissue or cell, which resulted in 256 total ramp sequence calculations for each variant (62 tissues + 66 cell types for both the reference and mutant sequences). By default, ExtRamp identifies ramp sequences based on codon translational efficiencies spanning nine codons, which is roughly the size of a ribosome window^61^. The harmonic mean is then used to determine the translational rate within that window, which is compared to the harmonic mean translational efficiency of the entire gene sequence. True outlier regions that occur at the 5’ end of genes are considered ramp sequences and were reported for each tissue and cell type. All scripts used to identify ramp sequences are available at https://github.com/jmillerlab/PILRA_ramp.

### Biological Assessment of Ramp Sequence Effects in PILRA

Synonymous variant *rs2405442:T>C* in *PILRA* was the only AD-associated variant predicted to destroy a ramp sequence. Since all five *PILRA* isoforms are predicted to have ramp sequences, we opted to use the longest *PILRA* isoform (Ensembl accession: *ENST00000198536.7*; NCBI accession: *NM_013439.3*) to assess *rs2405442:T>C* effects on mRNA and protein levels. DNA sequences for *ENST00000198536.7* (wildtype) and *ENST00000198536.7* containing *rs2405442:T>C* (mutant) were synthesized by GenScript Biotech. A Human c-Myc proto-oncogene (*MYC*) epitope tag, FLAG® epitope tag, and enterokinase cleavage site were attached to the 3’ end of the coding sequences. The reference and mutant sequences with annotated features are depicted in Supplementary Figures S1-S2.

Three independent replicates of quantitative polymerase chain reactions (qPCR) that each contained eight technical replicates were used to assess how synonymous variant, *rs2405442:T>C*, impacted *PILRA* mRNA levels in both the mutant and the wildtype transfected cells. Similarly, three independent sets of eight technical replicates were used to assess how PILRA protein levels differed between the mutant and wildtype using Enzyme-Linked Immunosorbent Assay (ELISA). Detailed instructions for replicating each protocol are described below.

### Transfection of wildtype and mutant transcripts

The wildtype and mutant sequences synthesized by GenScript Biotech were each inserted into separate mammalian expression vectors pCMV6-AN-myc-DDK (ORIGENE, Catalog #PS100016). Transformation protocols were followed as recommended by the manufacturer. In brief, plasmids were transformed into competent DH5a cells, amplified, and purified using the ZymoPURE II Plasmid Maxiprep kit (Catalog #D4203). Purified wildtype and mutant plasmids were then transfected into Chinese Hamster Ovary-K1 (CHO-K1) cells using Lipofectamine™ 3000 Transfection Reagent protocol (ThermoFisher, Catalog #15338100). Properly transfected cells were selected using the antibiotic G418 sulfate (ThermoFisher, Catalog #10131035). Transfected cells were grown in F12 media (ThemoFisher, Catalog #11765054) with 10 mg/ml penicillin, 10 μg/ml streptomycin (Gibco, Catalog #21127-022), and 10% FBS (HYCLone Catalog #SH30071.01). Cell media was changed every 48 to 72 hours depending on cell confluency levels. CHO-K1 transfected cells were then used in both the qPCR and ELISA protocols.

### qPCR Protocol

Total RNA was extracted from the mutant and the wildtype CHO-K1 transfected cells using the SPLIT RNA Extraction Kit (Lexogen, Catalog #008) and following the manufacturer guidelines. When the total RNA was purified and ready for quality control, the total RNA concentration was quantified using a NanoDrop spectrophotometer, and the Agilent DNF-471 RNA Kit 15nt (Agilent, Catalog #DNF-471-0500). Reverse transcription of the RNA into complementary DNA (cDNA) was then performed to convert the RNA molecules into their corresponding cDNA sequences using the High-Capacity cDNA Reverse Transcription Kit (Thermo Fisher Scientific, Catalog #4374966). We then performed qPCR to quantify gene expression levels, and cDNA concentration was quantified using the Agilent Femto Pulse. *PILRA* mRNA expression was normalized to total RNA to account for potential differences in qPCR amplification efficiency between tests. We calculated the relative expression of *PILRA* (ΔC_t_) by subtracting the *PILRA* counts (C_t_) from a housekeeping gene, Glyceraldehyde 3-phosphate dehydrogenase (*GAPDH*; i.e., *PILRA* – *GAPDH*). True outliers were then removed to limit potential technical artifacts. Finally, we calculated the fold change in expression (ΔΔC_t_)^62^ using the following equation with the average ΔC_t MUTANT_ and ΔC_t CONTROL_ across all replicates: ΔΔC_t_ = 2^−(ΔCt^ ^MUTANT−ΔCt^ ^CONTROL)^

### ELISA Protocol

Proteins were extracted from the mutant and the wildtype CHO-K1 transfected cells, and the total protein concentration was quantified using the Pierce™ BCA Protein Assay (Thermo Fisher Scientific, Catalog #23225 and 23227) following the manufacturer guidelines. The human PILRA protein concentration was quantitively measured in the mutant and the wildtype CHO-K1 transfected cells using the Human PILR-alpha ELISA Kit (Thermo Fisher Scientific, Catalog #EH368RB) by following the manufacturer guidelines. The concentration of PILRA was then normalized to the total concentration of proteins to account for potential variation between tests.

## RESULTS

### Ramp Sequence Variation Caused by Exonic GWAS Hits

Table 1 lists the 14 credibly causal exonic SNPs spanning 12 genes and 79 isoforms reported by Jansen, et al. ^54^. Each SNP was previously associated with AD and is here reported with the following: CADD^56^ score from GRCh38-v1.6; highest MAF reported in Ensembl^55^ from any population in 1000G Phase 3^63^, NHLBI Exome Sequencing Project^64^, and gnomAD^65^; GERP^57^ score from 91_mammals.gerp_conservation_score; RegulomeDB^58^ score; and effect on ramp sequences. Six SNPs (*rs2405442:T>C*, *rs12453:T>C*, *rs7982:A>G*, *rs1859788:A>G*, *rs12459419:C>T*, and *rs2296160:A>G*) are in five genes (*PILRA*, *MS4A6A*, *CR1*, *CLU*, and *CD33*) with ramp sequences. While *rs2405442:T>C*, *rs12453:T>C*, *rs1859788:A>G*, and *rs12459419:C>T* change the ramp sequence length, only *rs2405442:T>C* has a severe impact on ramp sequences by destroying it in at least one tissue or cell type.

**Table 1:**
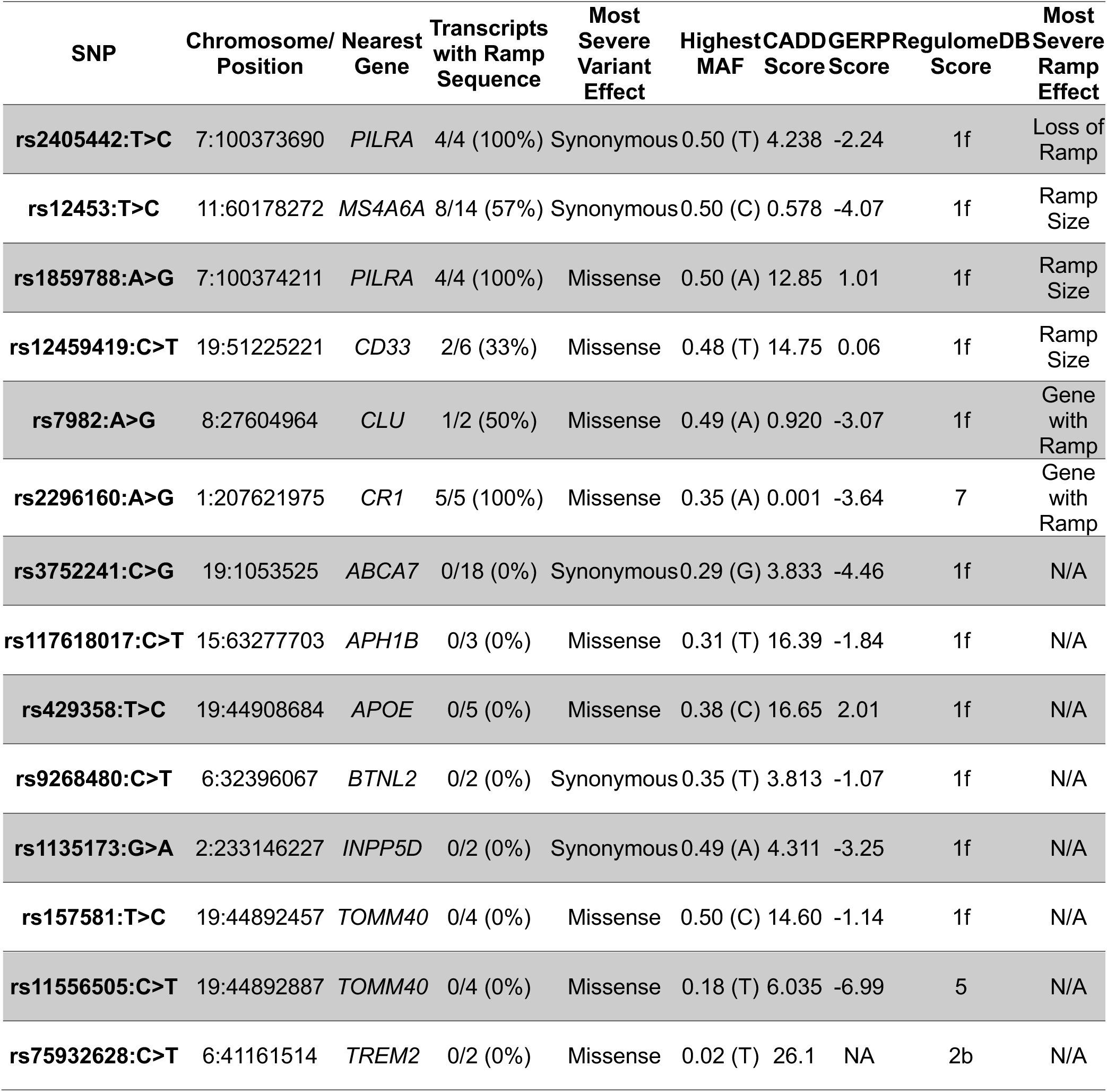
Credible Causal Exonic Variant Effects. Credible causality is defined and reported by Jansen, et al. ^54^. “Loss of Ramp” indicates that the ramp sequence was destroyed in at least one transcript, while “Ramp Size” indicates that the length of the ramp sequence changed in at least one transcript. “Gene with Ramp” indicates that the SNP was located outside the ramp region, yet the gene has a ramp sequence in at least one transcript in at least one cell or tissue.

### PILRA Ramp Sequence

Using ExtRamp, we calculated the relative codon adaptiveness of *PILRA* using all four isoforms in GRCh38. We then calculated the relative codon adaptiveness of *PILRA* with synonymous variant *rs2405442:T>C* and found that a ramp sequence is present in the reference isoforms but not the mutant isoforms for all four transcripts. The *PILRA* SNP, *rs2405442:T>C*, increases the regional mean translational efficiency at the 5’ end of *PILRA*, effectively destroying the ramp sequence (see Figure 1).

**Figure 1:**
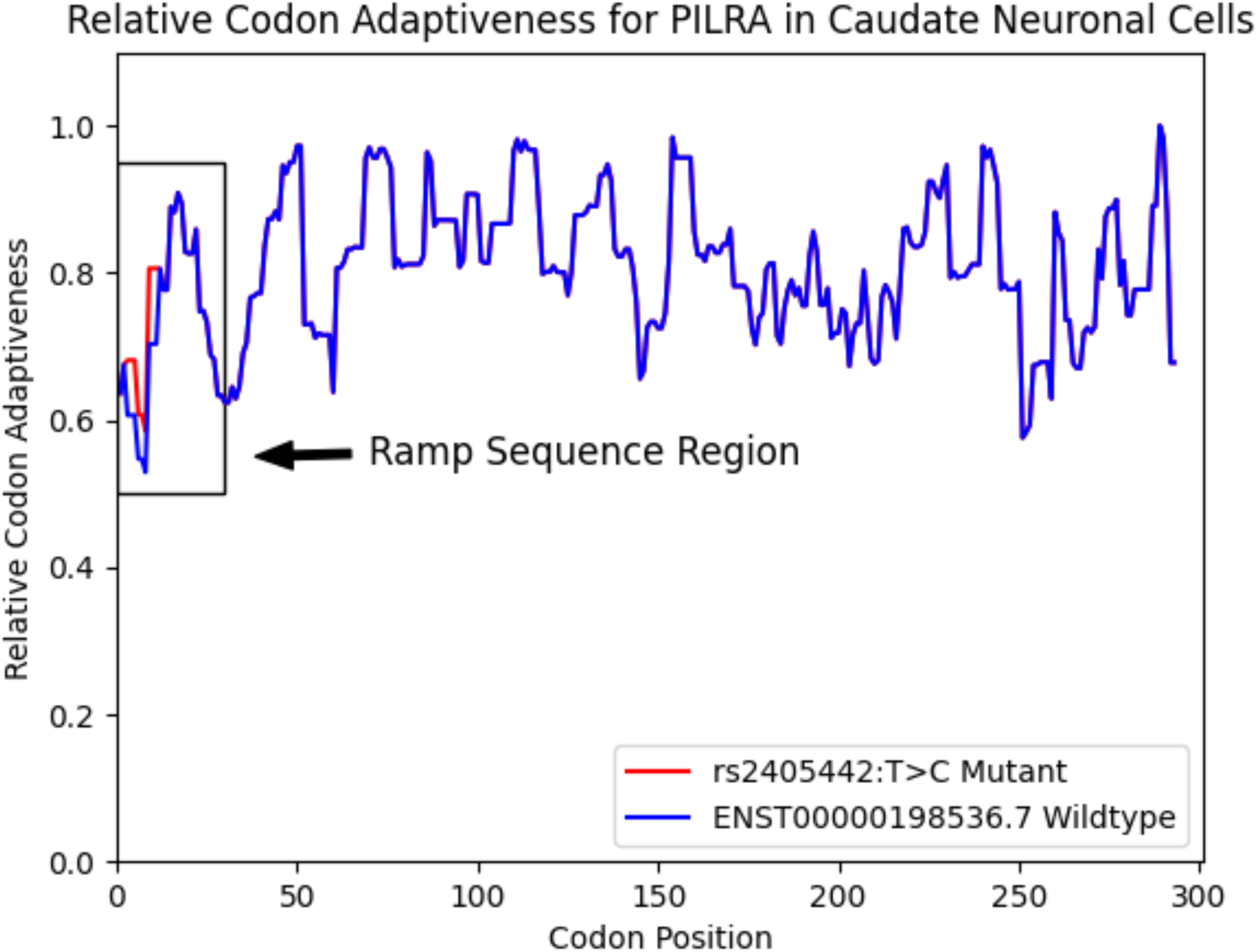
Figure 1 shows the relative codon adaptiveness of the longest PILRA reference isoform and the mutant gene averaged over a nine-codon window in caudate neuronal cells. The mutant gene (*rs2405442:T>C*) has a higher codon adaptiveness at the beginning of the gene sequence compared to the wildtype gene.

Using the consensus dataset consisting of gene expression from GTEx, FANTOM5, and the Human Protein Atlas, *PILRA* has a ramp sequence in 26/62 tissues. Using a single-cell dataset from the Human Protein Atlas, we also predicted that ramp sequences occur in *PILRA* in 20/66 cell types. Synonymous variant *rs2405442:T>C* destroyed the ramp sequence in all 46 tissues and cell types that normally contain a ramp sequence. Table 2 lists the 46 tissues or cell types with ramp sequences in *PILRA* that are affected by *rs2405442:T>C* (see Supplementary Table S1 for tissues and cell types without a *PILRA* ramp sequence). Specific neural cells that lost their ramp sequence include cerebellum Purkinje, hippocampus glial, caudate glial, and caudate neuronal cells. Lymphatic tissues and cells that lost their ramp sequence include the dendritic cells, monocytes, appendix lymphoid tissue, lymph node non-germinal center cells, and spleen cells in the red and white pulps.

**Table 2:**
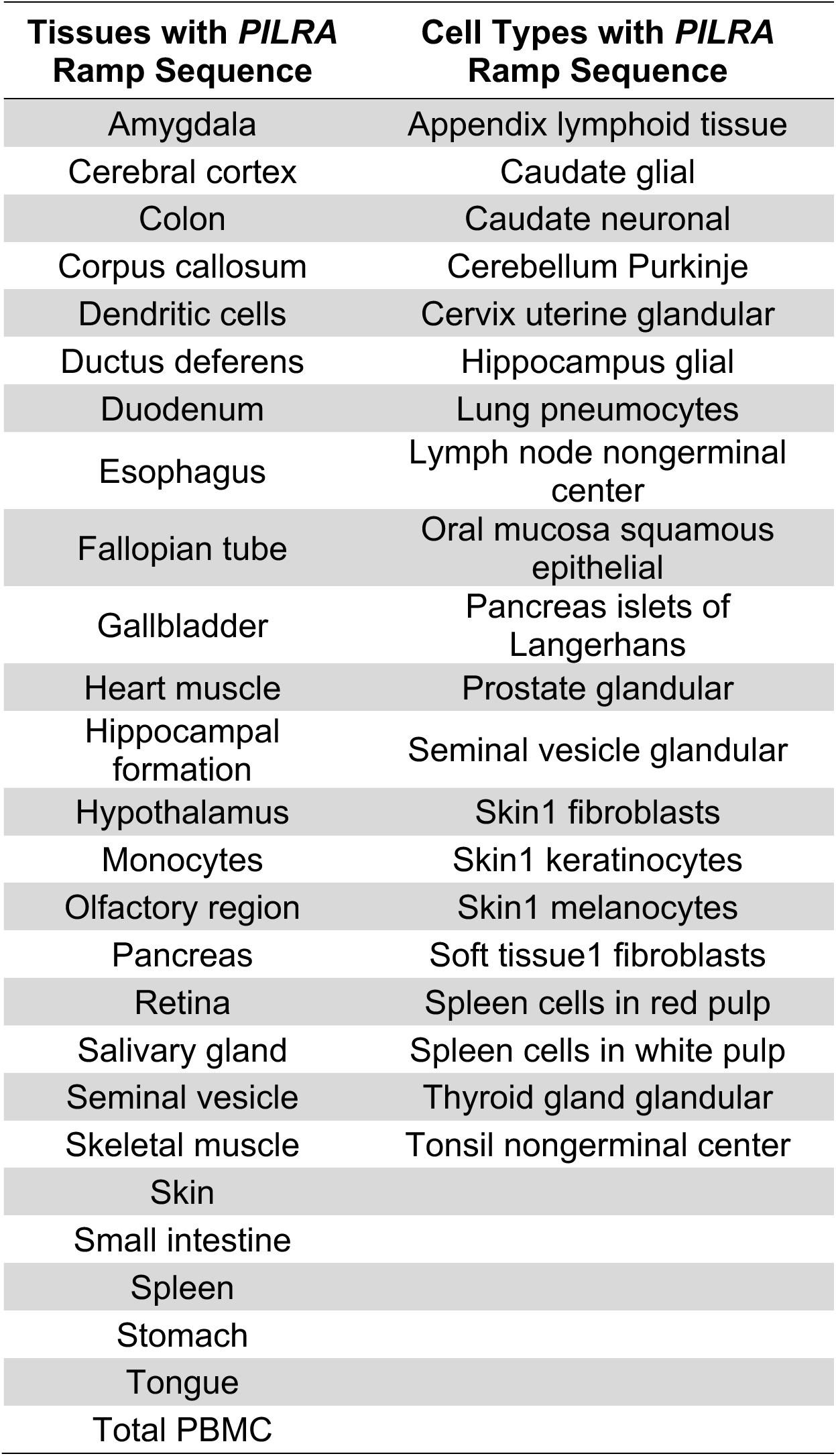
Tissues and cell types that normally have a ramp sequence in *PILRA*. The ramp sequence is universally destroyed in the mutant containing *rs2405442:T>C*.

In addition to ramp sequences, we also evaluated the effects of *rs2405442:T>C* on other codon usage biases such as GC content, codon pairing^66^, codon aversion^67–70^, and codon translational speed. *PILRA* was not previously identified as having splicing quantitative trait loci (sQTLs)^71^, so *rs2405442:T>C* is not predicted to impact splicing. GC content is slightly increased in the mutant, which would normally indicate higher mRNA expression^72^. Additionally, *rs2405442:T>C* affects the twelfth codon in *PILRA*, changing it from an uncommon leucine-encoding codon, TTG, to the most common leucine-encoding codon, CTG, which would normally indicate a higher translational speed since common codons are generally translated faster than rare codons^73^. Similarly, identical codon pairing suggests that *rs2405442:T>C* would increase translational speed^66^ since the mutation increases CTG codon pairing in the transcript from 6 instances in the wildtype to 7 instances in the mutant. Since the synonymous variant does not change the amino acid sequence, co-tRNA codon pairings (i.e., co-occurring amino acid residues)^74^ were not assessed. Based on codon usage biases, the ramp sequence indicates decreased mRNA and protein expression while GC content and codon pairing would suggest increased mRNA and protein expression in the mutant versus the wildtype.

Since synonymous variant *rs2405442:T>C* is the only credible causal synonymous variant with a predicted deleterious effect on a ramp sequence and is highly associated with AD, it was a good candidate for biological validation. We predicted that the destroyed ramp sequence would have an outsized effect due to ribosome-associated protein quality control induced by increased ribosome collisions^41^, and we experimentally validated the predicted effects of *rs2405442:T>C* on *PILRA* mRNA and protein levels with qPCR and ELISA using CHO cells harboring the synonymous variant compared to wildtype cells without the variant.

Figure 2 shows that mRNA levels are significantly lower in the mutant than the wildtype (*P*=1.9184 × 10^−13^). The fold change in expression (ΔΔC_t_) is ~131x higher in the wildtype than the mutant. Similarly, protein levels are also significantly higher in the wildtype than the mutant (*P*=0.01296), with *PILRA* protein levels being, on average, 1.1635x higher in the wildtype cells than the mutant cells. Although synonymous variant *rs2405442:T>C* has no effect on *PILRA* amino acid residues, it significantly decreases both mRNA and protein levels in the mutant versus the wildtype.

**Figure 2:**
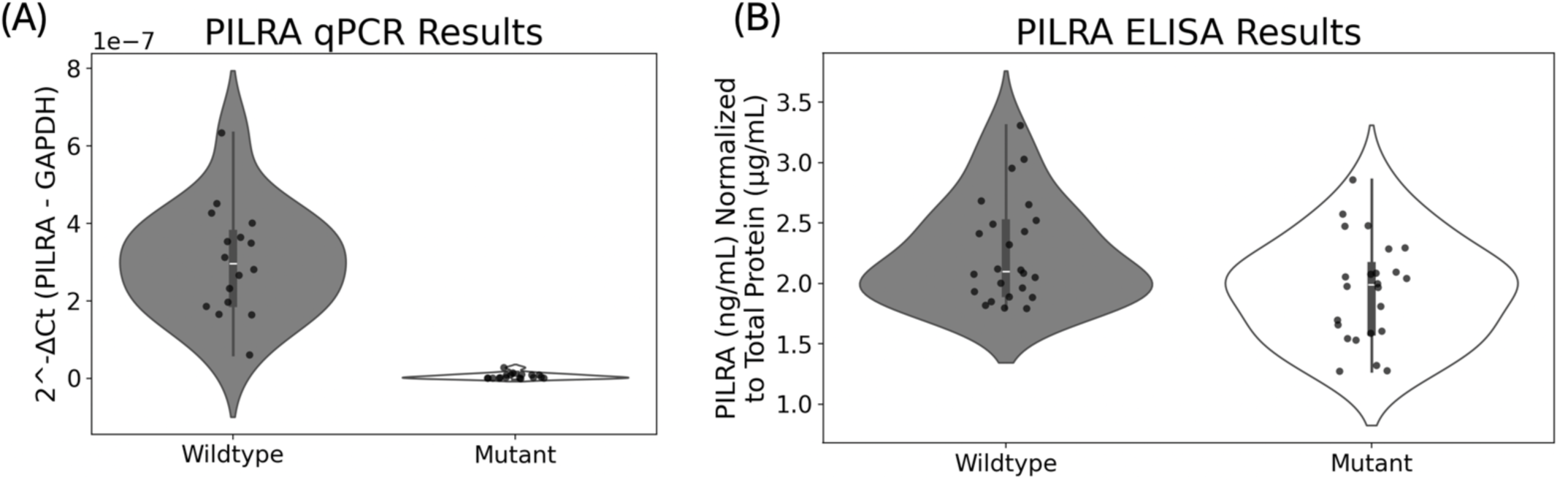
*rs2405442:T>C* effects on mRNA and protein levels in CHO cells harboring the synonymous variant compared to wildtype cells without the variant. (A) shows that *PILRA* mRNA levels are significantly lower in the mutant than the wildtype (P=1.9184 × 10^−13^). Since high C_t_ values show lower expression, we converted the raw C_t_ values to the relative expression by using the formula 2^−Ct^, where C_t_ is the normalized expression of C*_PILRA_* – C*_GAPDH_.* Two outliers with higher expression from the wildtype were removed, but did not affect the conclusions; (B) shows that *PILRA* protein levels are also significantly lower in the mutant than the wildtype (*P*=0.01296).

## DISCUSSION

Here, we provide a mechanistic explanation for the association of *rs2405442:T>C* with AD, including experimental validation of its biological effects. This study is the first time that ramp sequences have been used to prioritize disease-associated variants for biological validation. Since reduced *PILRA* inhibitory signaling has previously been shown to induce a protective effect against AD via reduced inhibitory signaling in microglia^53^, it is likely that less *PILRA* expression induced by *rs2405442:T>C* would similarly reduce inhibition of immune cells and result in more efficient cell-mediated clearance of Aβ. Although *rs2405442:T>C* creates a more common codon that increases GC content and codon pairing, which would generally increase mRNA and protein expression, the destroyed ramp sequence outweighs the other codon usage biases producing the observed effects. The destruction of the ramp sequence increases the frequency of ribosomal collisions, leading to stalled proteins and triggering the recruitment of the ribosome-associated protein quality control^41,75^ to degrade aberrant *PILRA* peptides.

*PILRA* gene expression *in vivo* is also likely affected by the distribution of tissue-specific mature tRNA pools. Some evidence suggests that mature tRNA pools change with environmental factors such as aging, stress, and diet, which would also change the relative codon adaptiveness and presence of a ramp sequence^76^. We show that *rs2405442:T>C* alone can significantly affect mRNA and protein levels independent of other genetic variants or changes in the tRNA pool, and further provide a workflow to perform tissue and cell specific computational analyses to investigate how a genetic variant impact gene-specific ramp sequences in different tissues and cells based on tRNA pool availability. Our tissue and cell-specific data show that after acquiring *rs2405442:T>C*, neural tissues such as caudate glial and neuronal cells, cerebellum Purkinje cells, and hippocampus glial cell are predicted to similarly lose their ramp sequence. Many lymphatic and immune-related tissues and cells likewise lose the *PILRA* ramp sequence after acquiring the synonymous variant, including dendritic cells, monocytes, PBMCs, appendix lymphoid tissue, lymph node non-germinal center cells, spleen cells in the red and white pulp, and tonsil non-germinal center cells. Since *PILRA* ramp sequence loss occurs in tissues and cell types known to impact AD, synonymous variant *rs2405442:T>C* in *PILRA* likely protects individuals from AD risk via reduced inhibitory signaling in those tissues and cells.

Synonymous variant *rs2405442:T>C* has a high minor allele frequency (MAF > 0.35^63,65^), indicating that natural decreases in *PILRA* expression induced by this variant are well-tolerated in the general population. Therefore, ramp-mediated therapeutics targeting *rs2405442:T>C* may be viable methods to mitigate risk for AD and increase cell-mediated Aβ clearance without inducing other off-target effects.

Here, we show that a ramp sequence plays a crucial role in *PILRA* gene regulation, and synonymous variant *rs2405442:T>C* alone causes a significant decrease in *PILRA m*RNA and protein levels by disrupting that regulatory mechanism. While synonymous variants are often overlooked in genome-wide association studies, they can significantly alter regulatory biases such as ramp sequences that can directly impact gene expression and protein levels. We outline how to analyze variant effects on ramp sequences, and we present a code repository at https://github.com/jmillerlab/PILRA_ramp to facilitate these types of analyses.

## ACKNOWLEDGEMENTS

We appreciate the contributions of the University of Kentucky Sanders-Brown Center on Aging and Brigham Young University for providing the research space and resources to conduct these analyses. We acknowledge the Office of Research Computing at Brigham Young University and the Center for Computational Sciences at the University of Kentucky for providing computational infrastructure and technical support. We also thank the University of Kentucky Center for Computational Sciences and Information Technology Services Research Computing for their support and use of the Morgan Compute Cluster and associated research computing resources.

## Data Availability

All scripts used for these analyses are publicly available at https://github.com/jmillerlab/PILRA_ramp.

## REFERENCES

1 Gatz, M. et al. Role of Genes and Environments for Explaining Alzheimer Disease. Archives of General Psychiatry 63, 168–174 (2006). 10.1001/archpsyc.63.2.168

2 Bellenguez, C. et al. New insights into the genetic etiology of Alzheimer’s disease and related dementias. Nature Genetics 54, 412–436 (2022). 10.1038/s41588-022-01024-z

3 Balin, B. J. & Hudson, A. P. Etiology and Pathogenesis of Late-Onset Alzheimer’s Disease. Current Allergy and Asthma Reports 14, 417 (2014). 10.1007/s11882-013-0417-1

4 Marioni, R. E. et al. GWAS on family history of Alzheimer’s disease. Translational Psychiatry 8, 99 (2018). 10.1038/s41398-018-0150-6

5 Jun, G. et al. Meta-analysis Confirms CR1, CLU, and PICALM as Alzheimer Disease Risk Loci and Reveals Interactions With APOE Genotypes. Archives of Neurology 67, 1473–1484 (2010). 10.1001/archneurol.2010.201

6 Hu, X. et al. Meta-Analysis for Genome-Wide Association Study Identifies Multiple Variants at the BIN1 Locus Associated with Late-Onset Alzheimer’s Disease. PLoS One 6 (2011). 10.1371/journal.pone.0016616

7 Lambert, J. C. et al. Meta-analysis of 74,046 individuals identifies 11 new susceptibility loci for Alzheimer’s disease. Nat Genet 45, 1452–1458 (2013). 10.1038/ng.2802

8 Ridge, P. G. et al. Assessment of the genetic variance of late-onset Alzheimer’s disease. Neurobiol Aging 41, 200.e213–200.e220 (2016). 10.1016/j.neurobiolaging.2016.02.024

9 Andrews, S. J. et al. The complex genetic architecture of Alzheimer’s disease: novel insights and future directions. EBioMedicine 90, 104511 (2023). 10.1016/j.ebiom.2023.104511

10 Escott-Price, V. et al. Common polygenic variation enhances risk prediction for Alzheimer’s disease. Brain 138, 3673–3684 (2015). 10.1093/brain/awv268

11 Bakulski, K. M. et al. Cumulative Genetic Risk and APOE ε4 Are Independently Associated With Dementia Status in a Multiethnic, Population-Based Cohort. Neurol Genet 7, e576 (2021). 10.1212/nxg.0000000000000576

12 Bredesen, D. E. Metabolic profiling distinguishes three subtypes of Alzheimer’s disease. Aging (Albany NY) 7, 595–600 (2015). 10.18632/aging.100801

13 Ferreira, D. et al. Distinct subtypes of Alzheimer’s disease based on patterns of brain atrophy: longitudinal trajectories and clinical applications. Sci Rep 7, 46263 (2017). 10.1038/srep46263

14 Eppig, J. S. et al. Statistically derived subtypes and associations with cerebrospinal fluid and genetic biomarkers in mild cognitive impairment: a latent profile analysis. Journal of the International Neuropsychological Society 23, 564–576 (2017).

15 Squitti, R. et al. Non-ceruloplasmin copper distincts subtypes in Alzheimer’s disease: a genetic study of ATP7B frequency. Molecular neurobiology 54, 671–681 (2017).

16 Mao, Y.-F., Guo, Z.-Y., Pu, J.-L., Chen, Y.-X. & Zhang, B.-R. Association of CD33 and MS4A cluster variants with Alzheimer’s disease in East Asian populations. Neuroscience letters 609, 235–239 (2015).

17 Mann, U. M., Mohr, E., Gearing, M. & Chase, T. N. Heterogeneity in Alzheimer’s disease: progression rate segregated by distinct neuropsychological and cerebral metabolic profiles. J Neurol Neurosurg Psychiatry 55, 956–959 (1992).

18 Na, H. K. et al. Malignant progression in parietal-dominant atrophy subtype of Alzheimer’s disease occurs independent of onset age. Neurobiol Aging 47, 149–156 (2016). 10.1016/j.neurobiolaging.2016.08.001

19 Park, J. Y. et al. Robust Identification of Alzheimer’s Disease subtypes based on cortical atrophy patterns. Sci Rep 7, 43270 (2017). 10.1038/srep43270

20 Persson, K. et al. MRI-assessed atrophy subtypes in Alzheimer’s disease and the cognitive reserve hypothesis. PLoS One 12, e0186595 (2017). 10.1371/journal.pone.0186595

21 Varol, E., Sotiras, A. & Davatzikos, C. HYDRA: Revealing heterogeneity of imaging and genetic patterns through a multiple max-margin discriminative analysis framework. NeuroImage 145, 346–364 (2017). 10.1016/j.neuroimage.2016.02.041

22 Mukherjee, S. et al. Genetic data and cognitively defined late-onset Alzheimer’s disease subgroups. Molecular Psychiatry (2018). 10.1038/s41380-018-0298-8

23 Warren, J. D., Fletcher, P. D. & Golden, H. L. The paradox of syndromic diversity in Alzheimer disease. Nat Rev Neurol 8, 451–464 (2012). 10.1038/nrneurol.2012.135

24 Jack, C. R., Jr., et al. A/T/N: An unbiased descriptive classification scheme for Alzheimer disease biomarkers. Neurology 87, 539–547 (2016). 10.1212/wnl.0000000000002923

25 Bondareff, W. et al. Age and histopathologic heterogeneity in Alzheimer’s disease: evidence for subtypes. Archives of general psychiatry 44, 412–417 (1987).

26 Crane, P. K. et al. Incidence of cognitively defined late-onset Alzheimer’s dementia subgroups from a prospective cohort study. Alzheimer’s & Dementia 13, 1307–1316 (2017). 10.1016/j.jalz.2017.04.011

27 Cummings, J. L. Cognitive and behavioral heterogeneity in Alzheimer’s disease: seeking the neurobiological basis. Neurobiology of aging 21, 845–861 (2000).

28 Larner, A. & Doran, M. Clinical phenotypic heterogeneity of Alzheimer’s disease associated with mutations of the presenilin–1 gene. Journal of neurology 253, 139–158 (2006).

29 Pillon, B., Dubois, B., Lhermitte, F. & Agid, Y. Heterogeneity of cognitive impairment in progressive supranuclear palsy, Parkinson’s disease, and Alzheimer’s disease. Neurology 36, 1179–1179 (1986).

30 Purcell, S. M. et al. Common polygenic variation contributes to risk of schizophrenia and bipolar disorder. Nature 460, 748–752 (2009). 10.1038/nature08185

31 Lewis, C. M. & Vassos, E. Polygenic risk scores: from research tools to clinical instruments. Genome Medicine 12, 44 (2020). 10.1186/s13073-020-00742-5

32 Dittmar, K. A., Goodenbour, J. M. & Pan, T. Tissue-specific differences in human transfer RNA expression. PLoS Genet 2, e221 (2006). 10.1371/journal.pgen.0020221

33 Waldman, Y. Y., Tuller, T., Shlomi, T., Sharan, R. & Ruppin, E. Translation efficiency in humans: tissue specificity, global optimization and differences between developmental stages. Nucleic Acids Res 38, 2964–2974 (2010). 10.1093/nar/gkq009

34 Tuller, T. & Zur, H. Multiple roles of the coding sequence 5’ end in gene expression regulation. Nucleic Acids Res 43, 13–28 (2015). 10.1093/nar/gku1313

35 Verma, M. et al. A short translational ramp determines the efficiency of protein synthesis. Nature Communications 10, 5774 (2019). 10.1038/s41467-019-13810-1

36 Miller, J. B., Brase, L. R. & Ridge, P. G. ExtRamp: a novel algorithm for extracting the ramp sequence based on the tRNA adaptation index or relative codon adaptiveness. Nucleic Acids Research 47, 1123–1131 (2019). 10.1093/nar/gky1193

37 Tuller, T. et al. Composite effects of gene determinants on the translation speed and density of ribosomes. Genome Biol 12, R110 (2011). 10.1186/gb-2011-12-11-r110

38 Tuller, T. et al. An Evolutionarily Conserved Mechanism for Controlling the Efficiency of Protein Translation. Cell 141, 344–354 (2010). 10.1016/j.cell.2010.03.031

39 Dana, A. & Tuller, T. The effect of tRNA levels on decoding times of mRNA codons. Nucleic Acids Res 42, 9171–9181 (2014). 10.1093/nar/gku646

40 Park, H. & Subramaniam, A. R. Inverted translational control of eukaryotic gene expression by ribosome collisions. PLoS Biol 17, e3000396 (2019). 10.1371/journal.pbio.3000396

41 Joazeiro, C. A. P. Mechanisms and functions of ribosome-associated protein quality control. Nature reviews. Molecular cell biology 20, 368–383 (2019). 10.1038/s41580-019-0118-2

42 McKinnon, L. M., Miller, J. B., Whiting, M. F., Kauwe, J. S. K. & Ridge, P. G. A comprehensive analysis of the phylogenetic signal in ramp sequences in 211 vertebrates. Sci Rep 11, 622 (2021). 10.1038/s41598-020-78803-3

43 Hodgman, M. W., Miller, J. B., Meurs, T. E. & Kauwe, J. S. K. CUBAP: an interactive web portal for analyzing codon usage biases across populations. Nucleic Acids Research 48, 11030–11039 (2020). 10.1093/nar/gkaa863

44 Miller, J. B. et al. The Ramp Atlas: facilitating tissue and cell-specific ramp sequence analyses through an intuitive web interface. NAR Genom Bioinform 4, lqac039 (2022). 10.1093/nargab/lqac039

45 Wang, J., Shiratori, I., Uehori, J., Ikawa, M. & Arase, H. Neutrophil infiltration during inflammation is regulated by PILRα via modulation of integrin activation. Nature Immunology 14, 34–40 (2013). 10.1038/ni.2456

46 Uhlén, M. et al. Tissue-based map of the human proteome. Science 347, 1260419 (2015). doi:10.1126/science.1260419

47 Pontén, F., Jirström, K. & Uhlen, M. The Human Protein Atlas—a tool for pathology. The Journal of Pathology 216, 387–393 (2008). 10.1002/path.2440

48 Kohyama, M. et al. Monocyte infiltration into obese and fibrilized tissues is regulated by PILRα. European Journal of Immunology 46, 1214–1223 (2016). 10.1002/eji.201545897

49 Selkoe, D. J. & Hardy, J. The amyloid hypothesis of Alzheimer’s disease at 25&#xa0;years. EMBO Molecular Medicine 8, 595–608-608 (2016). 10.15252/emmm.201606210

50 Huang, K.-L., et al. A common haplotype lowers PU.1 expression in myeloid cells and delays onset of Alzheimer’s disease. Nature Neuroscience 20, 1052–1061 (2017). 10.1038/nn.4587

51 Li, Y. et al. Genomics of Alzheimer’s disease implicates the innate and adaptive immune systems. Cell Mol Life Sci 78, 7397–7426 (2021). 10.1007/s00018-021-03986-5

52 Patel, T. et al. Whole-exome sequencing of the BDR cohort: evidence to support the role of the PILRA gene in Alzheimer’s disease. Neuropathology and Applied Neurobiology 44, 506–521 (2018). 10.1111/nan.12452

53 Rathore, N. et al. Paired Immunoglobulin-like Type 2 Receptor Alpha G78R variant alters ligand binding and confers protection to Alzheimer’s disease. PLoS Genet 14, e1007427 (2018). 10.1371/journal.pgen.1007427

54 Jansen, I. E. et al. Genome-wide meta-analysis identifies new loci and functional pathways influencing Alzheimer’s disease risk. Nature Genetics 51, 404–413 (2019). 10.1038/s41588-018-0311-9

55 Harrison, P. W. et al. Ensembl 2024. Nucleic Acids Research 52, D891–D899 (2024).

56 Rentzsch, P., Witten, D., Cooper, G. M., Shendure, J. & Kircher, M. CADD: predicting the deleteriousness of variants throughout the human genome. Nucleic Acids Research 47, D886–D894 (2019). 10.1093/nar/gky1016

57 Davydov, E. V. et al. Identifying a high fraction of the human genome to be under selective constraint using GERP++. PLoS computational biology 6, e1001025 (2010).

58 Dong, S. et al. Annotating and prioritizing human non-coding variants with RegulomeDB v. 2. Nature Genetics, 1–3 (2023).

59 Consortium, G. The GTEx Consortium atlas of genetic regulatory effects across human tissues. Science 369, 1318–1330 (2020).

60 Noguchi, S. et al. FANTOM5 CAGE profiles of human and mouse samples. Scientific data 4, 1–10 (2017).

61 Ingolia, N. T., Ghaemmaghami, S., Newman, J. R. & Weissman, J. S. Genome-wide analysis in vivo of translation with nucleotide resolution using ribosome profiling. Science 324, 218–223 (2009). 10.1126/science.1168978

62 Livak, K. J. & Schmittgen, T. D. Analysis of relative gene expression data using real-time quantitative PCR and the 2(-Delta Delta C(T)) Method. Methods 25, 402–408 (2001). 10.1006/meth.2001.1262

63 Clarke, L. et al. The 1000 Genomes Project: data management and community access. Nat Methods 9, 459–462 (2012). 10.1038/nmeth.1974

64 Auer, P. L. et al. Guidelines for Large-Scale Sequence-Based Complex Trait Association Studies: Lessons Learned from the NHLBI Exome Sequencing Project. Am J Hum Genet 99, 791–801 (2016). 10.1016/j.ajhg.2016.08.012

65 Chen, S. et al. A genome-wide mutational constraint map quantified from variation in 76,156 human genomes. bioRxiv, 2022.2003.2020.485034 (2022). 10.1101/2022.03.20.485034

66 Miller, J. B., McKinnon, L. M., Whiting, M. F., Kauwe, J. S. K. & Ridge, P. G. Codon Pairs are Phylogenetically Conserved: A comprehensive analysis of codon pairing conservation across the Tree of Life. PLoS One 15, e0232260 (2020). 10.1371/journal.pone.0232260

67 Miller, J. B., McKinnon, L. M., Whiting, M. F. & Ridge, P. G. Codon use and aversion is largely phylogenetically conserved across the tree of life. Mol Phylogenet Evol 144, 106697 (2020). 10.1016/j.ympev.2019.106697

68 Hodgman, M. W., Miller, J. B., Meurs, T. E. & Kauwe, J. S. K. CUBAP: an interactive web portal for analyzing codon usage biases across populations. Nucleic Acids Res 48, 11030–11039 (2020). 10.1093/nar/gkaa863

69 Miller, J. B., McKinnon, L. M., Whiting, M. F. & Ridge, P. G. CAM: an alignment-free method to recover phylogenies using codon aversion motifs. PeerJ 7, e6984 (2019). 10.7717/peerj.6984

70 Miller, J. B., Hippen, A. A., Belyeu, J. R., Whiting, M. F. & Ridge, P. G. Missing something? Codon aversion as a new character system in phylogenetics. Cladistics 33, 545–556 (2017). 10.1111/cla.12183

71 Yamaguchi, K. et al. Splicing QTL analysis focusing on coding sequences reveals mechanisms for disease susceptibility loci. Nature Communications 13, 4659 (2022). 10.1038/s41467-022-32358-1

72 Kudla, G., Lipinski, L., Caffin, F., Helwak, A. & Zylicz, M. High guanine and cytosine content increases mRNA levels in mammalian cells. PLoS Biol 4, e180 (2006). 10.1371/journal.pbio.0040180

73 Rodriguez, A., Wright, G., Emrich, S. & Clark, P. L. %MinMax: A versatile tool for calculating and comparing synonymous codon usage and its impact on protein folding. Protein Sci 27, 356–362 (2018). 10.1002/pro.3336

74 Cannarozzi, G. et al. A role for codon order in translation dynamics. Cell 141, 355–367 (2010). 10.1016/j.cell.2010.02.036

75 Brandman, O. & Hegde, R. S. Ribosome-associated protein quality control. Nat Struct Mol Biol 23, 7–15 (2016). 10.1038/nsmb.3147

76 Zhou, Z., Sun, B., Yu, D. & Bian, M. Roles of tRNA metabolism in aging and lifespan. Cell Death & Disease 12, 548 (2021). 10.1038/s41419-021-03838-x

