## Supplementary Information for "Ramp sequence may explain synonymous variant association with Alzheimer’s disease in the Paired Immunoglobulin-like Type 2 Receptor Alpha (*PILRA*)"

### Supplementary Tables

#### Supplementary Table S1: Tissues and Cell Types Without a Ramp Sequence in *PILRA*

| **Tissues without** *PILRA* Ramp Sequence | **Cell Types without** *PILRA* Ramp Sequence |
|---|---|
| Adipose tissue | Adrenal gland glandular cells |
| Adrenal gland | Appendix glandular cells |
| Appendix | Bone marrow hematopoietic |
| B-cells | Breast glandular |
| Basal ganglia | Breast myoepithelial |
| Bone marrow | Bronchus respiratory epithelial |
| Breast | Cerebellum glandular layer |
| Cerebellum | Cerebellum molecular layer |
| Cervix uterine | Cerebral cortex glial |
| Endometrium | Cerebral cortex neuronal |
| Epididymis | Cervix uterine squamous epithelial |
| Granulocytes | Colon endothelial |
| Kidney | Colon glandular |
| Liver | Colon peripheral nerve or ganglion |
| Lung | Duodenum glandular |
| Lymph node | Endometrium1 glandular |
| Midbrain | Endometrium2 glandular |
| NK-cells | Epididymis glandular |
| Ovary | Esophagus squamous epithelial |
| Parathyroid gland | Fallopian tube glandular |
| Pituitary gland | Gallbladder glandular |
| Placenta | Heart muscle myocytes |
| Pons and medulla | Hippocampus neuronal |
| Prostate | Kidney glomeruli |
| Rectum | Kidney tubules |
| Smooth muscle | Liver hepatocytes |

| | |
|---|---|
| Spinal cord | Lung macrophages |
| Substantia nigra | Lymph node germinal center |
| T-cells | Nasopharynx respiratory epithelial |
| Testis | Pancreas exocrine glandular |
| Thalamus | Parathyroid gland glandular |
| Thymus | Placenta decidual |
| Thyroid gland | Placenta trophoblastic |
| Tonsil | Rectum glandular |
| Urinary bladder | Salivary gland glandular |
| Vagina | Skin1 Langerhans |
| | Skin2 epidermal |
| | Small intestine glandular |
| | Stomach1 glandular |
| | Stomach2 glandular |
| | Testis cellsinseminiferousducts |
| | Testis Leydig |
| | Tonsil germinal center |
| | Tonsil squamous epithelial |
| | Urinary bladder urothelial |
| | Vagina squamous epithelial |

### Supplementary Figures

#### Supplementary Figure S1: Wildtype Sequence with Annotated Features

All features and annotations are displayed using SnapGene to illustrate the reference (wildtype) sequence that was transfected into CHO-K1 plasmids. The feature annotations comprise the following seven pages.

Sequence: PILRA_wt_Cterminaltag.dna (Circular / 5786 bp)
Enzymes: Unique 6+ Cutters (50 of 678 total)
Features: 18 total

Unique Cutters **Bold**

```
5'  AACAAAATATTAACGCTTACAATTTCCATTCGCCATTCAGGCTGCGCAACTGTTGGGAAGGGCGATCGGTGCGGGCCTCTTCGCT   85
3'  TTGTTTTATAATTGCGAATGTTAAAGGTAAGCGGTAAGTCCGACGCGTTGACAACCCTTCCCGCTAGCCACGCCCGGAGAAGCGA
    [f1 ori]

    ATTACGCCAGCTGGCGAAAGGGGGATGTGCTGCAAGGCGATTAAGTTGGGTAACGCCAGGGTTTTCCCAGTCACGACGTTGTAAA   170
    TAATGCGGTCGACCGCTTTCCCCCTACACGACGTTCCGCTAATTCAACCCATTGCGGTCCCAAAAGGGTCAGTGCTGCAACATTT
                                                                                  [M13 fwd]

    ACGACGGCCAGTGCCAAGCTGATCTATACATTGAATCAATATTGGCAATTAGCCATATTAGTCATTGGTTATATAGCATAAATCA   255
    TGCTGCCGGTCACGGTTCGACTAGATATGTAACTTAGTTATAACCGTTAATCGGTATAATCAGTAACCAATATATCGTATTTAGT
    [M13 fwd]

                                                       BsrGI
    ATATTGGCTATTGGCCATTGCATACGTTGTATCTATATCATAATATGTACATTTATATTGGCTCATGTCCAATATGACCGCCATG   340
    TATAACCGATAACCGGTAACGTATGCAACATAGATATAGTATTATACATGTAAATATAACCGAGTACAGGTTATACTGGCGGTAC

                     SpeI     AseI
    TTGACATTGATTATTGACTAGTTATTAATAGTAATCAATTACGGGGTCATTAGTTCATAGCCCATATATGGAGTTCCGCGTTACA   425
    AACTGTAACTAATAACTGATCAATAATTATCATTAGTTAATGCCCCAGTAATCAAGTATCGGGTATATACCTCAAGGCGCAATGT
      [CMV enhancer]

    TAACTTACGGTAAATGGCCCGCCTGGCTGACCGCCCAACGACCCCCGCCCATTGACGTCAATAATGACGTATGTTCCCATAGTAA   510
    ATTGAATGCCATTTACCGGGCGGACCGACTGGCGGGTTGCTGGGGGCGGGTAACTGCAGTTATTACTGCATACAAGGGTATCATT
    [CMV enhancer]

    CGCCAATAGGGACTTTCCATTGACGTCAATGGGTGGAGTATTTACGGTAAACTGCCCACTTGGCAGTACATCAAGTGTATCATAT   595
    GCGGTTATCCCTGAAAGGTAACTGCAGTTACCCACCTCATAAATGCCATTTGACGGGTGAACCGTCATGTAGTTCACATAGTATA
    [CMV enhancer]

    GCCAAGTCCGCCCCCTATTGACGTCAATGACGGTAAATGGCCCGCCTGGCATTATGCCCAGTACATGACCTTACGGGACTTTCCT   680
    CGGTTCAGGCGGGGGATAACTGCAGTTACTGCCATTTACCGGGCGGACCGTAATACGGGTCATGTACTGGAATGCCCTGAAAGGA
    [CMV enhancer]

                      SnaBI
    ACTTGGCAGTACATCTACGTATTAGTCATCGCTATTACCATGGTGATGCGGTTTTGGCAGTACACCAATGGGCGTGGATAGCGGT   765
    TGAACCGTCATGTAGATGCATAATCAGTAGCGATAATGGTACCACTACGCCAAAACCGTCATGTGGTTACCCGCACCTATCGCCA
    [CMV enhancer]                          [CMV promoter>

    TTGACTCACGGGGATTTCCAAGTCTCCACCCCATTGACGTCAATGGGAGTTTGTTTTGGCACCAAAATCAACGGGACTTTCCAAA   850
    AACTGAGTGCCCCTAAAGGTTCAGAGGTGGGGTAACTGCAGTTACCCTCAAACAAAACCGTGGTTTTAGTTGCCCTGAAAGGTTT
    [CMV promoter>

    ATGTCGTAATAACCCCGCCCCGTTGACGCAAATGGGCGGTAGGCGTGTACGGTGGGAGGTCTATATAAGCAGAGCTCGTTTAGTG   935
    TACAGCATTATTGGGGCGGGGCAACTGCGTTTACCCGCCATCCGCACATGCCACCCTCCAGATATATTCGTCTCGAGCAAATCAC
    [CMV promoter>
```

EcoRI SalI AccI BamHI Acc65I KpnI BglII

AACCGTCAGAATTTTGTAATACGACTCACTATAGGGCGGCCGGGAATTCGTCGACTGGATCCGGTACCGAGGAGATCTGCCGCCG 1020
TTGGCAGTCTTAAAACATTATGCTGAGTGATATCCCGCCGGCCCTTAAGCAGCTGACCTAGGCCATGGCTCCTCTAGACGGCGGC

T7 promoter

AsiSI

CGATCGCGCCACCatgggtcggcccctgctgctgcccctactgcccttgctgctgccgccagcatttctgcagcctagtggctcc 1105
GCTAGCGCGGTGGtacccagccggggacgacgacggggatgacgggaacgacgacggcggtcgtaaagacgtcggatcaccgagg

PILRA_wt

Kozak sequence

wt_ttg

Met Gly Arg Pro Leu Leu Leu Pro Leu Leu Pro Leu Leu Leu Pro Pro Ala Phe Leu Gln Pro Ser Gly Ser

acaggatctggtccaagctacctttatggggtcactcaaccaaaacacctctcagcctccatgggtggctctgtggaaatcccct 1190
tgtcctagaccaggttcgatggaaataccccagtgagttggttttgtggagagtcggaggtacccaccgagacacctttagggga

PILRA_wt

Thr Gly Ser Gly Pro Ser Tyr Leu Tyr Gly Val Thr Gln Pro Lys His Leu Ser Ala Ser Met Gly Gly Ser Val Glu Ile Pro

BmgBI

tctccttctattaccctgggagttagccacagctcccgacgtgagaatatcctggagacggggccacttccacaggcagtcctt 1275
agaggaagataatgggaccctcaatcggtgtcgagggctgcactcttataggacctctgccccggtgaaggtgtccgtcaggaa

PILRA_wt

Phe Ser Phe Tyr Tyr Pro Trp Glu Leu Ala Thr Ala Pro Asp Val Arg Ile Ser Trp Arg Arg Gly His Phe His Arg Gln Ser Phe

ctacagcacaaggccgccttccattcacaaggattatgtgaaccggctctttctgaactggacagagggtcagaagagcggcttc 1360
gatgtcgtgttccggcggaaggtaagtgttcctaatacacttggccgagaaagacttgacctgtctcccagtcttctcgccgaag

PILRA_wt

Tyr Ser Thr Arg Pro Pro Ser Ile His Lys Asp Tyr Val Asn Arg Leu Phe Leu Asn Trp Thr Glu Gly Gln Lys Ser Gly Phe

ctcaggatctccaacctgcagaagcaggaccagtctgtgtatttctgccgagttgagctggacacacggagctcagggaggcagc 1445
gagtcctagaggttggacgtcttcgtcctggtcagacacataaagacggctcaactcgacctgtgtgcctcgagtccctccgtcg

PILRA_wt

Leu Arg Ile Ser Asn Leu Gln Lys Gln Asp Gln Ser Val Tyr Phe Cys Arg Val Glu Leu Asp Thr Arg Ser Ser Gly Arg Gln

agtggcagtccatcgaggggaccaaactctccatcacccaggctgtcacgaccaccacccagaggcccagcagcatgactaccac 1530
tcaccgtcaggtagctcccctggtttgagaggtagtgggtccgacagtgctggtggtgggtctccgggtcgtcgtactgatggtg

PILRA_wt

Gln Trp Gln Ser Ile Glu Gly Thr Lys Leu Ser Ile Thr Gln Ala Val Thr Thr Thr Thr Gln Arg Pro Ser Ser Met Thr Thr Thr

ctggaggctcagtagcacaaccaccacaaccggcctcagggtcacacagggcaaacgacgctcagactcttggcacataagtctg 1615
gacctccgagtcatcgtgttggtggtgttggccggagtcccagtgtgtcccgtttgctgcgagtctgagaaccgtgtattcagac

PILRA_wt

Trp Arg Leu Ser Ser Thr Thr Thr Thr Thr Gly Leu Arg Val Thr Gln Gly Lys Arg Arg Ser Asp Ser Trp His Ile Ser Leu

gagactgctgtgggggtggcagtggctgtcactgtgctcggaatcatgattttgggactgatctgcctcctcaggtggaggagaa 1700
ctctgacgacacccccaccgtcaccgacagtgacacgagccttagtactaaaaccctgactagacggaggagtccacctcctctt

PILRA_wt

Glu Thr Ala Val Gly Val Ala Val Ala Val Thr Val Leu Gly Ile Met Ile Leu Gly Leu Ile Cys Leu Leu Arg Trp Arg Arg

ggaaaggtcagcagcggactaaagccacaaccccagccagggaacccttccaaaacacagaggagccatatgagaatatcaggaa 1785
cctttccagtcgtcgcctgatttcggtgttggggtcggtcccttgggaaggttttgtgtctcctcggtatactcttatagtcctt
PILRA_wt
Arg Lys Gly Gln Gln Arg Thr Lys Ala Thr Thr Pro Ala Arg Glu Pro Phe Gln Asn Thr Glu Glu Pro Tyr Glu Asn Ile Arg Asn

tgaaggacaaaatacagatcccaagctaaatcccaaggatgacggcatcgtctatgcttcccttgccctctccagctccacctca 1870
acttcctgttttatgtctagggttcgatttagggttcctactgccgtagcagatacgaagggaacgggagaggtcgaggtggagt
PILRA_wt
Glu Gly Gln Asn Thr Asp Pro Lys Leu Asn Pro Lys Asp Asp Gly Ile Val Tyr Ala Ser Leu Ala Leu Ser Ser Ser Thr Ser

NotI XhoI PspXI PaeR7I
cccagagcacctcccagccaccgtcccctcaagagccccagaacgagaccctgtactctgtcttaaaggcGCGGCCGCTCGAGC 1955
gggtctcgtggagggtcggtggcaggggagttctcggggtcttgctctgggacatgagacagaatttccgCGCCGGCGAGCTCG
PILRA_wt 1 Glu Myc
Pro Arg Ala Pro Pro Ser His Arg Pro Leu Lys Ser Pro Gln Asn Glu Thr Leu Tyr Ser Val Leu Lys Ala Arg Pro Leu Glu

EcoRV PmeI FseI
AGAAACTCATCTCAGAAGAGGATCTGGCAGCAAATGATATCCTGGATTACAAGGATGACGACGATAAGGTTTAAACGGCCGGCCG 2040
TCTTTGAGTAGAGTCTTCTCCTAGACCGTCGTTTACTATAGGACCTAATGTTCCTACTGCTGCTATTCCAAATTTGCCGGCCGGC
5 10 1 5
Gln Lys Leu Ile Ser Glu Glu Asp Leu Ala Ala Asn Asp Ile Leu Asp Tyr Lys Asp Asp Asp Asp Lys Val
Myc (in frame with Myc) FLAG
Gln Lys Leu Ile Ser Glu Glu Asp Leu Ala Ala Asn Asp Ile Leu Asp Tyr Lys Asp Asp Asp Asp Lys Val

SacII XmaI TspMI SmaI
CGGTCATAGCTGTTTCCTGAACAGATCCCGGGTGGCATCCCTGTGACCCCTCCCCAGTGCCTCTCCTGGCCCTGGAAGTTGCCAC 2125
GCCAGTATCGACAAAGGACTTGTCTAGGGCCCACCGTAGGGACACTGGGGAGGGGTCACGGAGAGGACCGGGACCTTCAACGGTG
M13 rev hGH poly(A) signal

AleI
TCCAGTGCCCACCAGCCTTGTCCTAATAAAATTAAGTTGCATCATTTTGTCTGACTAGGTGTCCTTCTATAATATTATGGGGTGG 2210
AGGTCACGGGTGGTCGGAACAGGATTATTTTAATTCAACGTAGTAAAACAGACTGATCCACAGGAAGATATTATAATACCCCACC
hGH poly(A) signal

BbsI
AGGGGGGTGGTATGGAGCAAGGGGCAAGTTGGGAAGACAACCTGTAGGGCCTGCGGGGTCTATTGGGAACCAAGCTGGAGTGCAG 2295
TCCCCCCACCATACCTCGTTCCCCGTTCAACCCTTCTGTTGGACATCCCGGACGCCCCAGATAACCCTTGGTTCGACCTCACGTC
hGH poly(A) signal

BsgI BbvCI
TGGCACAATCTTGGCTCACTGCAATCTCCGCCTCCTGGGTTCAAGCGATTCTCCTGCCTCAGCCTCCCGAGTTGTTGGGATTCCA 2380
ACCGTGTTAGAACCGAGTGACGTTAGAGGCGGAGGACCCAAGTTCGCTAAGAGGACGGAGTCGGAGGGCTCAACAACCCTAAGGT
hGH poly(A) signal

BlpI
GGCATGCATGACCAGGCTCAGCTAATTTTTGTTTTTTTGGTAGAGACGGGGTTTCACCATATTGGCCAGGCTGGTCTCCAACTCC 2465
CCGTACGTACTGGTCCGAGTCGATTAAAAACAAAAAAACCATCTCTGCCCCAAAGTGGTATAACCGGTCCGACCAGAGGTTGAGG
hGH poly(A) signal

BstXI
TAATCTCAGGTGATCTACCCACCTTGGCCTCCCAAATTGCTGGGATTACAGGCGTGAACCACTGCTCCCTTCCCTGTCCTTCTGA 2550
ATTAGAGTCCACTAGATGGGTGGAACCGGAGGGTTTAACGACCCTAATGTCCGCACTTGGTGACGAGGGAAGGGACAGGAAGACT
hGH poly(A) signal

AgeI PflMI
TTTTAAAATAACTATACCAGCAGGAGGACGTCCAGACACAGCATAGGCTACCTGGCCATGCCCAACCGGTGGGACATTTGAGTTG 2635
AAAATTTTATTGATATGGTCGTCCTCCTGCAGGTCTGTGTCGTATCCGATGGACCGGTACGGGTTGGCCACCCTGTAAACTCAAC
hGH poly(A) signal

CTTGCTTGGCACTGTCCTCTCATGCGTTGGGTCCACTCAGTAGATGCCTGTTGAATTGGGTACGCGGCCAGCGGCGAGCGGTATC 2720
GAACGAACCGTGACAGGAGAGTACGCAACCCAGGTGAGTCATCTACGGACAACTTAACCCATGCGCCGGTCGCCGCTCGCCATAG
hGH poly(A) signal

PciI
AflIII
AGCTCACTCAAAGGCGGTAATACGGTTATCCACAGAATCAGGGGATAACGCAGGAAAGAACATGTGAGCAAAAGGCCAGCAAAAG 2805
TCGAGTGAGTTTCCGCCATTATGCCAATAGGTGTCTTAGTCCCCTATTGCGTCCTTTCTTGTACACTCGTTTTCCGGTCGTTTTC

GCCAGGAACCGTAAAAAGGCCGCGTTGCTGGCGTTTTTCCATAGGCTCCGCCCCCCTGACGAGCATCACAAAAATCGACGCTCAA 2890
CGGTCCTTGGCATTTTTCCGGCGCAACGACCGCAAAAAGGTATCCGAGGCGGGGGGACTGCTCGTAGTGTTTTTAGCTGCGAGTT
ori

GTCAGAGGTGGCGAAACCCGACAGGACTATAAAGATACCAGGCGTTTCCCCCTGGAAGCTCCCTCGTGCGCTCTCCTGTTCCGAC 2975
CAGTCTCCACCGCTTTGGGCTGTCCTGATATTTCTATGGTCCGCAAAGGGGGACCTTCGAGGGAGCACGCGAGAGGACAAGGCTG
ori

CCTGCCGCTTACCGGATACCTGTCCGCCTTTCTCCCTTCGGGAAGCGTGGCGCTTTCTCATAGCTCACGCTGTAGGTATCTCAGT 3060
GGACGGCGAATGGCCTATGGACAGGCGGAAAGAGGGAAGCCCTTCGCACCGCGAAAGAGTATCGAGTGCGACATCCATAGAGTCA
ori

ApaLI
TCGGTGTAGGTCGTTCGCTCCAAGCTGGGCTGTGTGCACGAACCCCCCGTTCAGCCCGACCGCTGCGCCTTATCCGGTAACTATC 3145
AGCCACATCCAGCAAGCGAGGTTCGACCCGACACACGTGCTTGGGGGGCAAGTCGGGCTGGCGACGCGGAATAGGCCATTGATAG
ori

GTCTTGAGTCCAACCCGGTAAGACACGACTTATCGCCACTGGCAGCAGCCACTGGTAACAGGATTAGCAGAGCGAGGTATGTAGG 3230
CAGAACTCAGGTTGGGCCATTCTGTGCTGAATAGCGGTGACCGTCGTCGGTGACCATTGTCCTAATCGTCTCGCTCCATACATCC
ori

CGGTGCTACAGAGTTCTTGAAGTGGTGGCCTAACTACGGCTACACTAGAAGAACAGTATTTGGTATCTGCGCTCTGCTGAAGCCA 3315
GCCACGATGTCTCAAGAACTTCACCACCGGATTGATGCCGATGTGATCTTCTTGTCATAAACCATAGACGCGAGACGACTTCGGT
ori

GTTACCTTCGGAAAAAGAGTTGGTAGCTCTTGATCCGGCAAACAAACCACCGCTGGTAGCGGTGGTTTTTTTGTTTGCAAGCAGC 3400
CAATGGAAGCCTTTTTCTCAACCATCGAGAACTAGGCCGTTTGTTTGGTGGCGACCATCGCCACCAAAAAAACAAACGTTCGTCG
ori

```
AGATTACGCGCAGAAAAAAAGGATCTCAAGAAGATCCTTTGATCTTTTCTACGGGGTCTGACGCTCAGTGGAACGAAAACTCACG
TCTAATGCGCGTCTTTTTTTCCTAGAGTTCTTCTAGGAAACTAGAAAAGATGCCCCAGACTGCGAGTCACCTTGCTTTTGAGTGC   3485
ori

TTAAGGGATTTTGGTCATGAGATTATCAAAAAGGATCTTCACCTAGATCCTTTTAAATTAAAAATGAAGTTTTAAATCAATCTAA
AATTCCCTAAAACCAGTACTCTAATAGTTTTTCCTAGAAGTGGATCTAGGAAAATTTAATTTTTACTTCAAAATTTAGTTAGATT   3570

AGTATATATGAGTAACCTGAGGCTATGGCAGGGCCTGCCGCCCCGACGTTGGCTGCGAGCCCTGGGCCTTCACCCGAACTTGGGG
TCATATATACTCATTGGACTCCGATACCGTCCCGGACGGCGGGGCTGCAACCGACGCTCGGGACCCGGAAGTGGGCTTGAACCCC   3655

GGTGGGGTGGGGAAAAGGAAGAAACGCGGGCGTATTGGCCCCAATGGGGTCTCGGTGGGGTATCGACAGAGTGCCAGCCCTGGGA
CCACCCCACCCCTTTTCCTTCTTTGCGCCCGCATAACCGGGGTTACCCCAGAGCCACCCCATAGCTGTCTCACGGTCGGGACCCT   3740

CCGAACCCCGCGTTTATGAACAAACGACCCAACACCGTGCGTTTTATTCTGTCTTTTTATTGCCGTCATAGCGCGGGTTCCTTCC
GGCTTGGGGCGCAAATACTTGTTTGCTGGGTTGTGGCACGCAAAATAAGACAGAAAAATAACGGCAGTATCGCGCCCAAGGAAGG   3825
HSV TK poly(A) signal

GGTATTGTCTCCTTCCGTGTTTCAGTTAGCCTCCCCCTAGGGTGGGCGAAGAACTCCAGCATGAGATCCCCGCGCTGGAGGATCA
CCATAACAGAGGAAGGCACAAAGTCAATCGGAGGGGGATCCCACCCGCTTCTTGAGGTCGTACTCTAGGGGCGCGACCTCCTAGT   3910

TCCAGCCGGCGTCCCGGAAAACGATTCCGAAGCCCAACCTTTCATAGAAGGCGGCGGTGGAATCGAAATCTCGTGATGGCAGGTT
AGGTCGGCCGCAGGGCCTTTTGCTAAGGCTTCGGGTTGGAAAGTATCTTCCGCCGCCACCTTAGCTTTAGAGCACTACCGTCCAA   3995

BstBI
GGGCGTCGCTTGGTCGGTCATTTCGAACCCCAGAGTCCCGCTCAGAAGAACTCGTCAAGAAGGCGATAGAAGGCGATGCGCTGCG
CCCGCAGCGAACCAGCCAGTAAAGCTTGGGGTCTCAGGGCGAGTCTTCTTGAGCAGTTCTTCCGCTATCTTCCGCTACGCGACGC   4080
265 260 255
* Phe Phe Glu Asp Leu Leu Arg Tyr Phe Ala Ile Arg Gln Ser
NeoR/KanR
* Phe Phe Glu Asp Leu Leu Arg Tyr Phe Ala Ile Arg Gln Ser

AATCGGGAGCGGCGATACCGTAAAGCACGAGGAAGCGGTCAGCCCATTCGCCGCCAAGCTCTTCAGCAATATCACGGGTAGCCAA
TTAGCCCTCGCCGCTATGGCATTTCGTGCTCCTTCGCCAGTCGGGTAAGCGGCGGTTCGAGAAGTCGTTATAGTGCCCATCGGTT   4165
250 245 240 235 230 225
Asp Pro Ala Ala Ile Gly Tyr Leu Val Leu Phe Arg Asp Ala Trp Glu Gly Gly Leu Glu Glu Ala Ile Asp Arg Thr Ala Leu
NeoR/KanR
Asp Pro Ala Ala Ile Gly Tyr Leu Val Leu Phe Arg Asp Ala Trp Glu Gly Gly Leu Glu Glu Ala Ile Asp Arg Thr Ala Leu

CGCTATGTCCTGATAGCGATCCGCCACACCCAGCCGGCCACAGTCGATGAATCCAGAAAAGCGGCCATTTTCCACCATGATATTC
GCGATACAGGACTATCGCTAGGCGGTGTGGGTCGGCCGGTGTCAGCTACTTAGGTCTTTTCGCCGGTAAAAGGTGGTACTATAAG   4250
220 215 210 205 200 195
Ala Ile Asp Gln Tyr Arg Asp Ala Val Gly Leu Arg Gly Cys Asp Ile Phe Gly Ser Phe Arg Gly Asn Glu Val Met Ile Asn
NeoR/KanR
Ala Ile Asp Gln Tyr Arg Asp Ala Val Gly Leu Arg Gly Cys Asp Ile Phe Gly Ser Phe Arg Gly Asn Glu Val Met Ile Asn

GGCAAGCAGGCATCGCCATGGGTCACGACGAGATCCTCGCCGTCGGGCATGCTCGCCTTGAGCCTGGCGAACAGTTCGGCTGGCG
CCGTTCGTCCGTAGCGGTACCCAGTGCTGCTCTAGGAGCGGCAGCCCGTACGAGCGGAACTCGGACCGCTTGTCAAGCCGACCGC   4335
190 185 180 175 170
Pro Leu Cys Ala Asp Gly His Thr Val Val Leu Asp Glu Gly Asp Pro Met Ser Ala Lys Leu Arg Ala Phe Leu Glu Ala Pro Ala
NeoR/KanR
Pro Leu Cys Ala Asp Gly His Thr Val Val Leu Asp Glu Gly Asp Pro Met Ser Ala Lys Leu Arg Ala Phe Leu Glu Ala Pro Ala
```

```
CGAGCCCCTGATGCTCTTCGTCCAGATCATCCTGATCGACAAGACCGGCTTCCATCCGAGTACGTGCTCGCTCGATGCGATGTTT   4420
GCTCGGGGACTACGAGAAGCAGGTCTAGTAGGACTAGCTGTTCTGGCCGAAGGTAGGCTCATGCACGAGCGAGCTACGCTACAAA
165              160              155              150              145              140
Leu Gly Gln His Glu Glu Asp Leu Asp Asp Gln Asp Val Leu Gly Ala Glu Met Arg Thr Arg Ala Arg Glu Ile Arg His Lys
< NeoR/KanR
Leu Gly Gln His Glu Glu Asp Leu Asp Asp Gln Asp Val Leu Gly Ala Glu Met Arg Thr Arg Ala Arg Glu Ile Arg His Lys
```

```
CGCTTGGTGGTCGAATGGGCAGGTAGCCGGATCAAGCGTATGCAGCCGCCGCATTGCATCAGCCATGATGGATACTTTCTCGGCA   4505
GCGAACCACCAGCTTACCCGTCCATCGGCCTAGTTCGCATACGTCGGCGGCGTAACGTAGTCGGTACTACCTATGAAAGAGCCGT
      135              130              125              120              115              110
Ala Gln His Asp Phe Pro Cys Thr Ala Pro Asp Leu Thr His Leu Arg Arg Met Ala Asp Ala Met Ile Ser Val Lys Glu Ala
< NeoR/KanR
Ala Gln His Asp Phe Pro Cys Thr Ala Pro Asp Leu Thr His Leu Arg Arg Met Ala Asp Ala Met Ile Ser Val Lys Glu Ala
```

```
                                                                            PflFI
                                                                            Tth111I
GGAGCAAGGTGAGATGACAGGAGATCCTGCCCCGGCACTTCGCCCAATAGCAGCCAGTCCCTTCCCGCTTCAGTGACAACGTCGA   4590
CCTCGTTCCACTCTACTGTCCTCTAGGACGGGGCCGTGAAGCGGGTTATCGTCGGTCAGGGAAGGGCGAAGTCACTGTTGCAGCT
            105              100              95               90               85
Pro Ala Leu His Ser Ser Leu Leu Asp Gln Gly Pro Val Glu Gly Leu Leu Leu Trp Asp Arg Gly Ala Glu Thr Val Val Asp Leu
< NeoR/KanR
Pro Ala Leu His Ser Ser Leu Leu Asp Gln Gly Pro Val Glu Gly Leu Leu Leu Trp Asp Arg Gly Ala Glu Thr Val Val Asp Leu
```

```
GCACAGCTGCGCAAGGAACGCCCGTCGTGGCCAGCCACGATAGCCGCGCTGCCTCGTCTTGCAGTTCATTCAGGGCACCGGACAG   4675
CGTGTCGACGCGTTCCTTGCGGGCAGCACCGGTCGGTGCTATCGGCGCGACGGAGCAGAACGTCAAGTAAGTCCCGTGGCCTGTC
80               75               70               65               60               55
Val Ala Ala Cys Pro Val Gly Thr Thr Ala Leu Trp Ser Leu Arg Ala Ala Glu Asp Gln Leu Glu Asn Leu Ala Gly Ser Leu
< NeoR/KanR
Val Ala Ala Cys Pro Val Gly Thr Thr Ala Leu Trp Ser Leu Arg Ala Ala Glu Asp Gln Leu Glu Asn Leu Ala Gly Ser Leu
```

```
                    NarI SfoI
               KasI           PluTI
GTCGGTCTTGACAAAAAGAACCGGGCGCCCTGCGCTGACAGCCGGAACACGGCGGCATCAGAGCAGCCGATTGTCTGTTGTGCC   4760
CAGCCAGAACTGTTTTTCTTGGCCCGCGGGACGCGACTGTCGGCCTTGTGCCGCCGTAGTCTCGTCGGCTAACAGACAACACGG
      50               45               40               35               30               25
Asp Thr Lys Val Phe Leu Val Pro Arg Gly Gln Ala Ser Leu Arg Phe Val Ala Ala Asp Ser Cys Gly Ile Thr Gln Gln Ala
< NeoR/KanR
Asp Thr Lys Val Phe Leu Val Pro Arg Gly Gln Ala Ser Leu Arg Phe Val Ala Ala Asp Ser Cys Gly Ile Thr Gln Gln Ala
```

```
CAGTCATAGCCGAATAGCCTCTCCACCCAAGCGGCCGGAGAACCTGCGTGCAATCCATCTTGTTCAATCATGCGAAACGATCCTC   4845
GTCAGTATCGGCTTATCGGAGAGGTGGGTTCGCCGGCCTCTTGGACGCACGTTAGGTAGAACAAGTTAGTACGCTTTGCTAGGAG
            20               15               10               5            1
Trp Asp Tyr Gly Phe Leu Arg Glu Val Trp Ala Ala Pro Ser Gly Ala His Leu Gly Asp Gln Glu Ile Met
< NeoR/KanR
Trp Asp Tyr Gly Phe Leu Arg Glu Val Trp Ala Ala Pro Ser Gly Ala His Leu Gly Asp Gln Glu Ile Met
```

```
           ClaI*
           BspDI*                StuI                                               SfiI
ATCCTGTCTCTTGATCGATCTTTGCAAAAGCCTAGGCCTCCAAAAAAGCCTCCTCACTACTTCTGGAATAGCTCAGAGGCCGAGG   4930
TAGGACAGAGAACTAGCTAGAAACGTTTTCGGATCCGGAGGTTTTTTCGGAGGAGTGATGAAGACCTTATCGAGTCTCCGGCTCC
              < SV40 promoter
                         SV40 ori
```

```
CGGCCTCGGCCTCTGCATAAATAAAAAAAATTAGTCAGCCATGGGGCGGAGAATGGGCGGAACTGGGCGGAGTTAGGGGCGGGAT   5015
GCCGGAGCCGGAGACGTATTTATTTTTTTTAATCAGTCGGTACCCCGCCTCTTACCCGCCTTGACCCGCCTCAATCCCCGCCCTA
< SV40 promoter
SV40 ori

GGGCGGAGTTAGGGGCGGGACTATGGTTGCTGACTAATTGAGATGCATGCTTTGCATACTTCTGCCTGCTGGGGAGCCTGGGGAC   5100
CCCGCCTCAATCCCCGCCCTGATACCAACGACTGATTAACTCTACGTACGAAACGTATGAAGACGGACGACCCCTCGGACCCCTG
< SV40 promoter

         CsiI
         SexAI*
TTTCCACACCTGGTTGCTGACTAATTGAGATGCATGCTTTGCATACTTCTGCCTGCTGGGGAGCCTGGGGACTTTCCACACCCTA   5185
AAAGGTGTGGACCAACGACTGATTAACTCTACGTACGAAACGTATGAAGACGGACGACCCCTCGGACCCCTGAAAGGTGTGGGAT
< SV40 promoter

ACTGACACACATTCCACAGCTGGTTCTTTCCGCCTCAGGACTCTTCCTTTTTCAATATTATTGAAGCATTTATCAGGGTTATTGT   5270
TGACTGTGTGTAAGGTGTCGACCAAGAAAGGCGGAGTCCTGAGAAGGAAAAAGTTATAATAACTTCGTAAATAGTCCCAATAACA
< SV40 promoter   < AmpR promoter

CTCATGAGCGGATACATATTTGAATGTATTTAGAAAAATAAACAAATAGGGGTTCCGCGCACATTTCCCCGAAAAGTGCCACCTG   5355
GAGTACTCGCCTATGTATAAACTTACATAAATCTTTTTATTTGTTTATCCCCAAGGCGCGTGTAAAGGGGCTTTTCACGGTGGAC
< AmpR promoter

ACGCGCCCTGTAGCGGCGCATTAAGCGCGGCGGGTGTGGTGGTTACGCGCAGCGTGACCGCTACACTTGCCAGCGCCCTAGCGCC   5440
TGCGCGGGACATCGCCGCGTAATTCGCGCCGCCCACACCACCAATGCGCGTCGCACTGGCGATGTGAACGGTCGCGGGATCGCGG
f1 ori >

CGCTCCTTTCGCTTTCTTCCCTTCCTTTCTCGCCACGTTCGCCGGCTTTCCCCGTCAAGCTCTAAATCGGGGGCTCCCTTTAGGG   5525
GCGAGGAAAGCGAAAGAAGGGAAGGAAAGAGCGGTGCAAGCGGCCGAAAGGGGCAGTTCGAGATTTAGCCCCCGAGGGAAATCCC
f1 ori >

                                                                 DraIII
TTCCGATTTAGTGCTTTACGGCACCTCGACCCCAAAAAACTTGATTAGGGTGATGGTTCACGTAGTGGGCCATCGCCCTGATAGA   5610
AAGGCTAAATCACGAAATGCCGTGGAGCTGGGGTTTTTTGAACTAATCCCACTACCAAGTGCATCACCCGGTAGCGGGACTATCT
f1 ori >

CGGTTTTTCGCCCTTTGACGTTGGAGTCCACGTTCTTTAATAGTGGACTCTTGTTCCAAACTGGAACAACACTCAACCCTATCTC   5695
GCCAAAAAGCGGGAAACTGCAACCTCAGGTGCAAGAAATTATCACCTGAGAACAAGGTTTGACCTTGTTGTGAGTTGGGATAGAG
f1 ori >

                   PsiI
GGTCTATTCTTTTGATTTATAAGGGATTTTGCCGATTTCGGCCTATTGGTTAAAAAATGAGCTGATTTAACAAAAATTTAACGCG   5780
CCAGATAAGAAAACTAAATATTCCCTAAAACGGCTAAAGCCGGATAACCAATTTTTTACTCGACTAAATTGTTTTTAAATTGCGC
f1 ori >

AATTTT   3'
         ...   5786
TTAAAA   5'
f1 ori >
```

#### Supplementary Figure S2: Mutant Sequence with Annotated Features

All features and annotations are displayed using SnapGene to illustrate the mutant sequence that was transfected into CHO-K1 plasmids. The only difference between the mutant and the wildtype sequence is *rs2405442:T>C*. Feature annotations comprise the following eight pages.

Sequence: PILRA_mt_Cterminaltag.dna (Circular / 5786 bp)
Enzymes: Unique 6+ Cutters (50 of 678 total)　　Unique Cutters **Bold**
Features: 18 total

```
5' AACAAAATATTAACGCTTACAATTTCCATTCGCCATTCAGGCTGCGCAACTGTTGGGAAGGGCGATCGGTGCGGGCCTCT
                                                                                        80
3' TTGTTTTATAATTGCGAATGTTAAAGGTAAGCGGTAAGTCCGACGCGTTGACAACCCTTCCCGCTAGCCACGCCCGGAGA
```

f1 ori

```
TCGCTATTACGCCAGCTGGCGAAAGGGGGATGTGCTGCAAGGCGATTAAGTTGGGTAACGCCAGGGTTTTCCCAGTCACG
                                                                                     160
AGCGATAATGCGGTCGACCGCTTTCCCCCTACACGACGTTCCGCTAATTCAACCCATTGCGGTCCCAAAAGGGTCAGTGC
```

```
ACGTTGTAAAACGACGGCCAGTGCCAAGCTGATCTATACATTGAATCAATATTGGCAATTAGCCATATTAGTCATTGGTT
                                                                                     240
TGCAACATTTTGCTGCCGGTCACGGTTCGACTAGATATGTAACTTAGTTATAACCGTTAATCGGTATAATCAGTAACCAA
```

M13 fwd

**BsrGI**

```
ATATAGCATAAATCAATATTGGCTATTGGCCATTGCATACGTTGTATCTATATCATAATATGTACATTTATATTGGCTCA
                                                                                     320
TATATCGTATTTAGTTATAACCGATAACCGGTAACGTATGCAACATAGATATAGTATTATACATGTAAATATAACCGAGT
```

**SpeI** **AseI**

```
TGTCCAATATGACCGCCATGTTGACATTGATTATTGACTAGTTATTAATAGTAATCAATTACGGGGTCATTAGTTCATAG
                                                                                     400
ACAGGTTATACTGGCGGTACAACTGTAACTAATAACTGATCAATAATTATCATTAGTTAATGCCCCAGTAATCAAGTATC
```

CMV enhancer

```
CCCATATATGGAGTTCCGCGTTACATAACTTACGGTAAATGGCCCGCCTGGCTGACCGCCCAACGACCCCCGCCCATTGA
                                                                                     480
GGGTATATACCTCAAGGCGCAATGTATTGAATGCCATTTACCGGGCGGACCGACTGGCGGGTTGCTGGGGGCGGGTAACT
```

CMV enhancer

```
CGTCAATAATGACGTATGTTCCCATAGTAACGCCAATAGGGACTTTCCATTGACGTCAATGGGTGGAGTATTTACGGTAA
                                                                                     560
GCAGTTATTACTGCATACAAGGGTATCATTGCGGTTATCCCTGAAAGGTAACTGCAGTTACCCACCTCATAAATGCCATT
```

CMV enhancer

```
ACTGCCCACTTGGCAGTACATCAAGTGTATCATATGCCAAGTCCGCCCCCTATTGACGTCAATGACGGTAAATGGCCCGC
                                                                                     640
TGACGGGTGAACCGTCATGTAGTTCACATAGTATACGGTTCAGGCGGGGGATAACTGCAGTTACTGCCATTTACCGGGCG
```

CMV enhancer

**SnaBI**

```
CTGGCATTATGCCCAGTACATGACCTTACGGGACTTTCCTACTTGGCAGTACATCTACGTATTAGTCATCGCTATTACCA
                                                                                     720
GACCGTAATACGGGTCATGTACTGGAATGCCCTGAAAGGATGAACCGTCATGTAGATGCATAATCAGTAGCGATAATGGT
```

CMV enhancer

```
TGGTGATGCGGTTTTGGCAGTACACCAATGGGCGTGGATAGCGGTTTGACTCACGGGGATTTCCAAGTCTCCACCCCATT
                                                                                     800
ACCACTACGCCAAAACCGTCATGTGGTTACCCGCACCTATCGCCAAACTGAGTGCCCCTAAAGGTTCAGAGGTGGGGTAA
```

CMV promoter

CMV enhancer

```
GACGTCAATGGGAGTTTGTTTTGGCACCAAAATCAACGGGACTTTCCAAAATGTCGTAATAACCCCGCCCCGTTGACGCA
                                                                                     880
CTGCAGTTACCCTCAAACAAAACCGTGGTTTTAGTTGCCCTGAAAGGTTTTACAGCATTATTGGGGCGGGGCAACTGCGT
```

CMV promoter

```
AATGGGCGGTAGGCGTGTACGGTGGGAGGTCTATATAAGCAGAGCTCGTTTAGTGAACCGTCAGAATTTTGTAATACGAC
                                                                                     960
TTACCCGCCATCCGCACATGCCACCCTCCAGATATATTCGTCTCGAGCAAATCACTTGGCAGTCTTAAAACATTATGCTG
```

CMV promoter　　T7 promoter

EcoRI SalI AccI BamHI Acc65I KpnI BglII AsiSI

TCACTATAGGGCGGCCGGGAATTCGTCGACTGGATCCGGTACCGAGGAGATCTGCCGCCGCGATCGCGCCACCatgggtc 1040

AGTGATATCCCGCCGGCCCTTAAGCAGCTGACCTAGGCCATGGCTCCTCTAGACGGCGGCGCTAGCGCGGTGGtacccag

T7 promoter

PILRA_mt

Kozak sequence

Met Gly

ggcccctgctgctgcccctactgcccctgctgctgccgccagcatttctgcagcctagtggctccacaggatctggtcca 1120

ccggggacgacgacggggatgacggggacgacgacggcggtcgtaaagacgtcggatcaccgaggtgtcctagaccaggt

PILRA_mt

mt_ctg

Arg Pro Leu Leu Leu Pro Leu Leu Pro Leu Leu Leu Pro Pro Ala Phe Leu Gln Pro Ser Gly Ser Thr Gly Ser Gly Pro

agctacctttatggggtcactcaaccaaaacacctctcagcctccatgggtggctctgtggaaatccccttctccttcta 1200

tcgatggaaataccccagtgagttggttttgtggagagtcggaggtacccaccgagacacctttaggggaagaggaagat

PILRA_mt

Ser Tyr Leu Tyr Gly Val Thr Gln Pro Lys His Leu Ser Ala Ser Met Gly Gly Ser Val Glu Ile Pro Phe Ser Phe Tyr

BmgBI

ttacccctgggagttagccacagctcccgacgtgagaatatcctggagacggggccacttccacaggcagtccttctaca 1280

aatggggaccctcaatcggtgtcgagggctgcactcttataggacctctgccccggtgaaggtgtccgtcaggaagatgt

PILRA_mt

Tyr Pro Trp Glu Leu Ala Thr Ala Pro Asp Val Arg Ile Ser Trp Arg Arg Gly His Phe His Arg Gln Ser Phe Tyr

gcacaaggccgccttccattcacaaggattatgtgaaccggctctttctgaactggacagagggtcagaagagcggcttc 1360

cgtgttccggcggaaggtaagtgttcctaatacacttggccgagaaagacttgacctgtctcccagtcttctcgccgaag

PILRA_mt

Ser Thr Arg Pro Pro Ser Ile His Lys Asp Tyr Val Asn Arg Leu Phe Leu Asn Trp Thr Glu Gly Gln Lys Ser Gly Phe

ctcaggatctccaacctgcagaagcaggaccagtctgtgtatttctgccgagttgagctggacacacggagctcagggag 1440

gagtcctagaggttggacgtcttcgtcctggtcagacacataaagacggctcaactcgacctgtgtgcctcgagtccctc

PILRA_mt

Leu Arg Ile Ser Asn Leu Gln Lys Gln Asp Gln Ser Val Tyr Phe Cys Arg Val Glu Leu Asp Thr Arg Ser Ser Gly Arg

gcagcagtggcagtccatcgaggggaccaaactctccatcacccaggctgtcacgaccaccacccagaggcccagcagca 1520

cgtcgtcaccgtcaggtagctcccctggtttgagaggtagtgggtccgacagtgctggtggtgggtctccgggtcgtcgt

PILRA_mt

Gln Gln Trp Gln Ser Ile Glu Gly Thr Lys Leu Ser Ile Thr Gln Ala Val Thr Thr Thr Thr Gln Arg Pro Ser Ser

tgactaccacctggaggctcagtagcacaaccaccacaaccggcctcagggtcacacagggcaaacgacgctcagactct 1600

actgatggtggacctccgagtcatcgtgttggtggtgttggccggagtcccagtgtgtcccgtttgctgcgagtctgaga

PILRA_mt

Met Thr Thr Thr Trp Arg Leu Ser Ser Thr Thr Thr Thr Thr Gly Leu Arg Val Thr Gln Gly Lys Arg Arg Ser Asp Ser

tggcacataagtctggagactgctgtgggggtggcagtggctgtcactgtgctcggaatcatgattttgggactgatctg 1680
accgtgtattcagacctctgacgacacccccaccgtcaccgacagtgacacgagccttagtactaaaaccctgactagac
PILRA_mt
Trp His Ile Ser Leu Glu Thr Ala Val Gly Val Ala Val Ala Val Thr Val Leu Gly Ile Met Ile Leu Gly Leu Ile Cys

cctcctcaggtggaggagaaggaaaggtcagcagcggactaaagccacaaccccagccagggaacccttccaaaacacag 1760
ggaggagtccacctcctcttcctttccagtcgtcgcctgatttcggtgttggggtcggtcccttgggaaggttttgtgtc
PILRA_mt
Leu Leu Arg Trp Arg Arg Arg Lys Gly Gln Gln Arg Thr Lys Ala Thr Thr Pro Ala Arg Glu Pro Phe Gln Asn Thr

aggagccatatgagaatatcaggaatgaaggacaaaatacagatcccaagctaaatcccaaggatgacggcatcgtctat 1840
tcctcggtatactcttatagtccttacttcctgttttatgtctagggttcgatttagggttcctactgccgtagcagata
PILRA_mt
Glu Glu Pro Tyr Glu Asn Ile Arg Asn Glu Gly Gln Asn Thr Asp Pro Lys Leu Asn Pro Lys Asp Asp Gly Ile Val Tyr

gcttcccttgccctctccagctccacctcacccagagcacctcccagccaccgtcccctcaagagccccccagaacgagac 1920
cgaagggaacgggagaggtcgaggtggagtgggtctcgtggagggtcggtggcaggggagttctcgggggtcttgctctg
PILRA_mt
Ala Ser Leu Ala Leu Ser Ser Ser Thr Ser Pro Arg Ala Pro Pro Ser His Arg Pro Leu Lys Ser Pro Gln Asn Glu Thr

XhoI
PspXI
NotI PaeR7I EcoRV
cctgtactctgtcttaaaggcGCGGCCGCTCGAGCAGAAACTCATCTCAGAAGAGGATCTGGCAGCAAATGATATCCTGG 2000
ggacatgagacagaatttccgCGCCGGCGAGCTCGTCTTTGAGTAGAGTCTTCTCCTAGACCGTCGTTTACTATAGGACC
1 5 10
Glu Gln Lys Leu Ile Ser Glu Glu Asp Leu Ala Ala Asn Asp Ile Leu
PILRA_mt
Myc
(in frame with Myc)
FLAG
Leu Tyr Ser Val Leu Lys Ala Arg Pro Leu Glu Gln Lys Leu Ile Ser Glu Glu Asp Leu Ala Ala Asn Asp Ile Leu

XmaI
PmeI FseI SacII TspMI SmaI
ATTACAAGGATGACGACGATAAGGTTTAAACGGCCGGCCGCGGTCATAGCTGTTTCCTGAACAGATCCCGGGTGGCATCC 2080
TAATGTTCCTACTGCTGCTATTCCAAATTTGCCGGCCGGCGCCAGTATCGACAAAGGACTTGTCTAGGGCCCACCGTAGG
1 5
Asp Tyr Lys Asp Asp Asp Asp Lys Val
FLAG
M13 rev
hGH poly(A) signal
Asp Tyr Lys Asp Asp Asp Asp Lys Val

AleI
CTGTGACCCCTCCCCAGTGCCTCTCCTGGCCCTGGAAGTTGCCACTCCAGTGCCCACCAGCCTTGTCCTAATAAAATTAA 2160
GACACTGGGGAGGGGTCACGGAGAGGACCGGGACCTTCAACGGTGAGGTCACGGGTGGTCGGAACAGGATTATTTTAATT
hGH poly(A) signal

GTTGCATCATTTTGTCTGACTAGGTGTCCTTCTATAATATTATGGGGTGGAGGGGGGTGGTATGGAGCAAGGGGCAAGTT 2240
CAACGTAGTAAAACAGACTGATCCACAGGAAGATATTATAATACCCCACCTCCCCCCACCATACCTCGTTCCCCGTTCAA
hGH poly(A) signal

BbsI, BsgI

GGGAAGACAACCTGTAGGGCCTGCGGGGTCTATTGGGAACCAAGCTGGAGTGCAGTGGCACAATCTTGGCTCACTGCAAT 2320
CCCTTCTGTTGGACATCCCGGACGCCCCAGATAACCCTTGGTTCGACCTCACGTCACCGTGTTAGAACCGAGTGACGTTA
hGH poly(A) signal

BbvCI, BlpI

CTCCGCCTCCTGGGTTCAAGCGATTCTCCTGCCTCAGCCTCCCGAGTTGTTGGGATTCCAGGCATGCATGACCAGGCTCA 2400
GAGGCGGAGGACCCAAGTTCGCTAAGAGGACGGAGTCGGAGGGCTCAACAACCCTAAGGTCCGTACGTACTGGTCCGAGT
hGH poly(A) signal

GCTAATTTTTGTTTTTTTGGTAGAGACGGGGTTTCACCATATTGGCCAGGCTGGTCTCCAACTCCTAATCTCAGGTGATC 2480
CGATTAAAAACAAAAAAACCATCTCTGCCCCAAAGTGGTATAACCGGTCCGACCAGAGGTTGAGGATTAGAGTCCACTAG
hGH poly(A) signal

BstXI

TACCCACCTTGGCCTCCCAAATTGCTGGGATTACAGGCGTGAACCACTGCTCCCTTCCCTGTCCTTCTGATTTTAAAATA 2560
ATGGGTGGAACCGGAGGGTTTAACGACCCTAATGTCCGCACTTGGTGACGAGGGAAGGGACAGGAAGACTAAAATTTTAT
hGH poly(A) signal

AgeI, PflMI

ACTATACCAGCAGGAGGACGTCCAGACACAGCATAGGCTACCTGGCCATGCCCAACCGGTGGGACATTTGAGTTGCTTGC 2640
TGATATGGTCGTCCTCCTGCAGGTCTGTGTCGTATCCGATGGACCGGTACGGGTTGGCCACCCTGTAAACTCAACGAACG
hGH poly(A) signal

TTGGCACTGTCCTCTCATGCGTTGGGTCCACTCAGTAGATGCCTGTTGAATTGGGTACGCGGCCAGCGGCGAGCGGTATC 2720
AACCGTGACAGGAGAGTACGCAACCCAGGTGAGTCATCTACGGACAACTTAACCCATGCGCCGGTCGCCGCTCGCCATAG
hGH poly(A) signal

PciI, AflIII

AGCTCACTCAAAGGCGGTAATACGGTTATCCACAGAATCAGGGGATAACGCAGGAAAGAACATGTGAGCAAAAGGCCAGC 2800
TCGAGTGAGTTTCCGCCATTATGCCAATAGGTGTCTTAGTCCCCTATTGCGTCCTTTCTTGTACACTCGTTTTCCGGTCG

AAAAGGCCAGGAACCGTAAAAAGGCCGCGTTGCTGGCGTTTTTCCATAGGCTCCGCCCCCCTGACGAGCATCACAAAAAT 2880
TTTTCCGGTCCTTGGCATTTTTCCGGCGCAACGACCGCAAAAAGGTATCCGAGGCGGGGGGACTGCTCGTAGTGTTTTTA
ori

CGACGCTCAAGTCAGAGGTGGCGAAACCCGACAGGACTATAAAGATACCAGGCGTTTCCCCCTGGAAGCTCCCTCGTGCG 2960
GCTGCGAGTTCAGTCTCCACCGCTTTGGGCTGTCCTGATATTTCTATGGTCCGCAAAGGGGGACCTTCGAGGGAGCACGC
ori

CTCTCCTGTTCCGACCCTGCCGCTTACCGGATACCTGTCCGCCTTTCTCCCTTCGGGAAGCGTGGCGCTTTCTCATAGCT 3040
GAGAGGACAAGGCTGGGACGGCGAATGGCCTATGGACAGGCGGAAAGAGGGAAGCCCTTCGCACCGCGAAAGAGTATCGA
ori

ApaLI

CACGCTGTAGGTATCTCAGTTCGGTGTAGGTCGTTCGCTCCAAGCTGGGCTGTGTGCACGAACCCCCCGTTCAGCCCGAC 3120
GTGCGACATCCATAGAGTCAAGCCACATCCAGCAAGCGAGGTTCGACCCGACACACGTGCTTGGGGGGCAAGTCGGGCTG
ori

CGCTGCGCCTTATCCGGTAACTATCGTCTTGAGTCCAACCCGGTAAGACACGACTTATCGCCACTGGCAGCAGCCACTGG 3200
GCGACGCGGAATAGGCCATTGATAGCAGAACTCAGGTTGGGCCATTCTGTGCTGAATAGCGGTGACCGTCGTCGGTGACC
ori

TAACAGGATTAGCAGAGCGAGGTATGTAGGCGGTGCTACAGAGTTCTTGAAGTGGTGGCCTAACTACGGCTACACTAGAA 3280
ATTGTCCTAATCGTCTCGCTCCATACATCCGCCACGATGTCTCAAGAACTTCACCACCGGATTGATGCCGATGTGATCTT
ori

GAACAGTATTTGGTATCTGCGCTCTGCTGAAGCCAGTTACCTTCGGAAAAAGAGTTGGTAGCTCTTGATCCGGCAAACAA 3360
CTTGTCATAAACCATAGACGCGAGACGACTTCGGTCAATGGAAGCCTTTTTCTCAACCATCGAGAACTAGGCCGTTTGTT
ori

ACCACCGCTGGTAGCGGTGGTTTTTTTGTTTGCAAGCAGCAGATTACGCGCAGAAAAAAAGGATCTCAAGAAGATCCTTT 3440
TGGTGGCGACCATCGCCACCAAAAAAACAAACGTTCGTCGTCTAATGCGCGTCTTTTTTTCCTAGAGTTCTTCTAGGAAA
ori

GATCTTTTCTACGGGGTCTGACGCTCAGTGGAACGAAAACTCACGTTAAGGGATTTTGGTCATGAGATTATCAAAAAGGA 3520
CTAGAAAAGATGCCCCAGACTGCGAGTCACCTTGCTTTTGAGTGCAATTCCCTAAAACCAGTACTCTAATAGTTTTTCCT

TCTTCACCTAGATCCTTTTAAATTAAAAATGAAGTTTTAAATCAATCTAAAGTATATATGAGTAACCTGAGGCTATGGCA 3600
AGAAGTGGATCTAGGAAAATTTAATTTTTACTTCAAAATTTAGTTAGATTTCATATATACTCATTGGACTCCGATACCGT

GGGCCTGCCGCCCCGACGTTGGCTGCGAGCCCTGGGCCTTCACCCGAACTTGGGGGGTGGGGTGGGGAAAAGGAAGAAAC 3680
CCCGGACGGCGGGGCTGCAACCGACGCTCGGGACCCGGAAGTGGGCTTGAACCCCCCACCCCACCCCTTTTCCTTCTTTG

GCGGGCGTATTGGCCCCAATGGGGTCTCGGTGGGGTATCGACAGAGTGCCAGCCCTGGGACCGAACCCCGCGTTTATGAA 3760
CGCCCGCATAACCGGGGTTACCCCAGAGCCACCCCATAGCTGTCTCACGGTCGGGACCCTGGCTTGGGGCGCAAATACTT
HSV TK poly(A) signal

CAAACGACCCAACACCGTGCGTTTTATTCTGTCTTTTTATTGCCGTCATAGCGCGGGTTCCTTCCGGTATTGTCTCCTTC 3840
GTTTGCTGGGTTGTGGCACGCAAAATAAGACAGAAAAATAACGGCAGTATCGCGCCCAAGGAAGGCCATAACAGAGGAAG
HSV TK poly(A) signal

CGTGTTTCAGTTAGCCTCCCCCTAGGGTGGGCGAAGAACTCCAGCATGAGATCCCCGCGCTGGAGGATCATCCAGCCGGC 3920
GCACAAAGTCAATCGGAGGGGGATCCCACCCGCTTCTTGAGGTCGTACTCTAGGGGCGCGACCTCCTAGTAGGTCGGCCG

GTCCCGGAAAACGATTCCGAAGCCCAACCTTTCATAGAAGGCGGCGGTGGAATCGAAATCTCGTGATGGCAGGTTGGGCG 4000
CAGGGCCTTTTGCTAAGGCTTCGGGTTGGAAAGTATCTTCCGCCGCCACCTTAGCTTTAGAGCACTACCGTCCAACCCGC

**BstBI**
TCGCTTGGTCGGTCATTTCGAACCCCAGAGTCCCGCTCAGAAGAACTCGTCAAGAAGGCGATAGAAGGCGATGCGCTGCG 4080
AGCGAACCAGCCAGTAAAGCTTGGGGTCTCAGGGCGAGTCTTCTTGAGCAGTTCTTCCGCTATCTTCCGCTACGCGACGC
265 260 255
* Phe Phe Glu Asp Leu Leu Arg Tyr Phe Ala Ile Arg Gln Ser
NeoR/KanR
* Phe Phe Glu Asp Leu Leu Arg Tyr Phe Ala Ile Arg Gln Ser

AATCGGGAGCGGCGATACCGTAAAGCACGAGGAAGCGGTCAGCCCATTCGCCGCCAAGCTCTTCAGCAATATCACGGGTA 4160
TTAGCCCTCGCCGCTATGGCATTTCGTGCTCCTTCGCCAGTCGGGTAAGCGGCGGTTCGAGAAGTCGTTATAGTGCCCAT

250 245 240 235 230 225
Asp Pro Ala Ala Ile Gly Tyr Leu Val Leu Phe Arg Asp Ala Trp Glu Gly Gly Leu Glu Glu Ala Ile Asp Arg Thr
NeoR/KanR
Asp Pro Ala Ala Ile Gly Tyr Leu Val Leu Phe Arg Asp Ala Trp Glu Gly Gly Leu Glu Glu Ala Ile Asp Arg Thr

GCCAACGCTATGTCCTGATAGCGATCCGCCACACCCAGCCGGCCACAGTCGATGAATCCAGAAAAGCGGCCATTTTCCAC 4240
CGGTTGCGATACAGGACTATCGCTAGGCGGTGTGGGTCGGCCGGTGTCAGCTACTTAGGTCTTTTCGCCGGTAAAAGGTG

220 215 210 205 200
Ala Leu Ala Ile Asp Gln Tyr Arg Asp Ala Val Gly Leu Arg Gly Cys Asp Ile Phe Gly Ser Phe Arg Gly Asn Glu Val
NeoR/KanR
Ala Leu Ala Ile Asp Gln Tyr Arg Asp Ala Val Gly Leu Arg Gly Cys Asp Ile Phe Gly Ser Phe Arg Gly Asn Glu Val

CATGATATTCGGCAAGCAGGCATCGCCATGGGTCACGACGAGATCCTCGCCGTCGGGCATGCTCGCCTTGAGCCTGGCGA 4320
GTACTATAAGCCGTTCGTCCGTAGCGGTACCCAGTGCTGCTCTAGGAGCGGCAGCCCGTACGAGCGGAACTCGGACCGCT

195 190 185 180 175
Met Ile Asn Pro Leu Cys Ala Asp Gly His Thr Val Val Leu Asp Glu Gly Asp Pro Met Ser Ala Lys Leu Arg Ala Phe
NeoR/KanR
Met Ile Asn Pro Leu Cys Ala Asp Gly His Thr Val Val Leu Asp Glu Gly Asp Pro Met Ser Ala Lys Leu Arg Ala Phe

ACAGTTCGGCTGGCGCGAGCCCCTGATGCTCTTCGTCCAGATCATCCTGATCGACAAGACCGGCTTCCATCCGAGTACGT 4400
TGTCAAGCCGACCGCGCTCGGGGACTACGAGAAGCAGGTCTAGTAGGACTAGCTGTTCTGGCCGAAGGTAGGCTCATGCA

170 165 160 155 150 145
Leu Glu Ala Pro Ala Leu Gly Gln His Glu Glu Asp Leu Asp Asp Gln Asp Val Leu Gly Ala Glu Met Arg Thr Arg
NeoR/KanR
Leu Glu Ala Pro Ala Leu Gly Gln His Glu Glu Asp Leu Asp Asp Gln Asp Val Leu Gly Ala Glu Met Arg Thr Arg

GCTCGCTCGATGCGATGTTTCGCTTGGTGGTCGAATGGGCAGGTAGCCGGATCAAGCGTATGCAGCCGCCGCATTGCATC 4480
CGAGCGAGCTACGCTACAAAGCGAACCACCAGCTTACCCGTCCATCGGCCTAGTTCGCATACGTCGGCGGCGTAACGTAG

140 135 130 125 120
Ala Arg Glu Ile Arg His Lys Ala Gln His Asp Phe Pro Cys Thr Ala Pro Asp Leu Thr His Leu Arg Arg Met Ala Asp
NeoR/KanR
Ala Arg Glu Ile Arg His Lys Ala Gln His Asp Phe Pro Cys Thr Ala Pro Asp Leu Thr His Leu Arg Arg Met Ala Asp

AGCCATGATGGATACTTTCTCGGCAGGAGCAAGGTGAGATGACAGGAGATCCTGCCCCGGCACTTCGCCCAATAGCAGCC 4560
TCGGTACTACCTATGAAAGAGCCGTCCTCGTTCCACTCTACTGTCCTCTAGGACGGGGCCGTGAAGCGGGTTATCGTCGG

115 110 105 100 95
Ala Met Ile Ser Val Lys Glu Ala Pro Ala Leu His Ser Ser Leu Leu Asp Gln Gly Pro Val Glu Gly Leu Leu Leu Trp
NeoR/KanR
Ala Met Ile Ser Val Lys Glu Ala Pro Ala Leu His Ser Ser Leu Leu Asp Gln Gly Pro Val Glu Gly Leu Leu Leu Trp

**PflFI**
**Tth111I**

AGTCCCTTCCCGCTTCAGTGACAACGTCGAGCACAGCTGCGCAAGGAACGCCCGTCGTGGCCAGCCACGATAGCCGCGCT 4640
TCAGGGAAGGGCGAAGTCACTGTTGCAGCTCGTGTCGACGCGTTCCTTGCGGGCAGCACCGGTCGGTGCTATCGGCGCGA

90 85 80 75 70 65
Asp Arg Gly Ala Glu Thr Val Val Asp Leu Val Ala Ala Cys Pro Val Gly Thr Thr Ala Leu Trp Ser Leu Arg Ala
NeoR/KanR
Asp Arg Gly Ala Glu Thr Val Val Asp Leu Val Ala Ala Cys Pro Val Gly Thr Thr Ala Leu Trp Ser Leu Arg Ala

KasI NarI SfoI PluTI

GCCTCGTCTTGCAGTTCATTCAGGGCACCGGACAGGTCGGTCTTGACAAAAAGAACCGGGCGCCCCTGCGCTGACAGCCG 4720
CGGAGCAGAACGTCAAGTAAGTCCCGTGGCCTGTCCAGCCAGAACTGTTTTTCTTGGCCCGCGGGGACGCGACTGTCGGC

60 55 50 45 40
Ala Glu Asp Gln Leu Glu Asn Leu Ala Gly Ser Leu Asp Thr Lys Val Phe Leu Val Pro Arg Gly Gln Ala Ser Leu Arg
NeoR/KanR
Ala Glu Asp Gln Leu Glu Asn Leu Ala Gly Ser Leu Asp Thr Lys Val Phe Leu Val Pro Arg Gly Gln Ala Ser Leu Arg

GAACACGGCGGCATCAGAGCAGCCGATTGTCTGTTGTGCCCAGTCATAGCCGAATAGCCTCTCCACCCAAGCGGCCGGAG 4800
CTTGTGCCGCCGTAGTCTCGTCGGCTAACAGACAACACGGGTCAGTATCGGCTTATCGGAGAGGTGGGTTCGCCGGCCTC

35 30 25 20 15
Phe Val Ala Ala Asp Ser Cys Gly Ile Thr Gln Gln Ala Trp Asp Tyr Gly Phe Leu Arg Glu Val Trp Ala Ala Pro Ser
NeoR/KanR
Phe Val Ala Ala Asp Ser Cys Gly Ile Thr Gln Gln Ala Trp Asp Tyr Gly Phe Leu Arg Glu Val Trp Ala Ala Pro Ser

ClaI* BspDI*

AACCTGCGTGCAATCCATCTTGTTCAATCATGCGAAACGATCCTCATCCTGTCTCTTGATCGATCTTTGCAAAAGCCTAG 4880
TTGGACGCACGTTAGGTAGAACAAGTTAGTACGCTTTGCTAGGAGTAGGACAGAGAACTAGCTAGAAACGTTTTCGGATC

10 5 1
Gly Ala His Leu Gly Asp Gln Glu Ile Met
NeoR/KanR
SV40 promoter
SV40 ori
Gly Ala His Leu Gly Asp Gln Glu Ile Met

StuI SfiI

GCCTCCAAAAAAGCCTCCTCACTACTTCTGGAATAGCTCAGAGGCCGAGGCGGCCTCGGCCTCTGCATAAATAAAAAAAA 4960
CGGAGGTTTTTTCGGAGGAGTGATGAAGACCTTATCGAGTCTCCGGCTCCGCCGGAGCCGGAGACGTATTTATTTTTTTT

SV40 promoter
SV40 ori

TTAGTCAGCCATGGGGCGGAGAATGGGCGGAACTGGGCGGAGTTAGGGGCGGGATGGGCGGAGTTAGGGGCGGGACTATG 5040
AATCAGTCGGTACCCCGCCTCTTACCCGCCTTGACCCGCCTCAATCCCCGCCCTACCCGCCTCAATCCCCGCCCTGATAC

SV40 promoter
SV40 ori

CsiI SexAI*

GTTGCTGACTAATTGAGATGCATGCTTTGCATACTTCTGCCTGCTGGGGAGCCTGGGGACTTTCCACACCTGGTTGCTGA 5120
CAACGACTGATTAACTCTACGTACGAAACGTATGAAGACGGACGACCCCTCGGACCCCTGAAAGGTGTGGACCAACGACT

SV40 promoter

CTAATTGAGATGCATGCTTTGCATACTTCTGCCTGCTGGGGAGCCTGGGGACTTTCCACACCCTAACTGACACACATTCC 5200
GATTAACTCTACGTACGAAACGTATGAAGACGGACGACCCCTCGGACCCCTGAAAGGTGTGGGATTGACTGTGTGTAAGG

SV40 promoter

ACAGCTGGTTCTTTCCGCCTCAGGACTCTTCCTTTTTCAATATTATTGAAGCATTTATCAGGGTTATTGTCTCATGAGCG 5280
TGTCGACCAAGAAAGGCGGAGTCCTGAGAAGGAAAAAGTTATAATAACTTCGTAAATAGTCCCAATAACAGAGTACTCGC

SV40 promoter AmpR promoter

GATACATATTTGAATGTATTTAGAAAAATAAACAAATAGGGGTTCCGCGCACATTTCCCCGAAAAGTGCCACCTGACGCG 5360
CTATGTATAAACTTACATAAATCTTTTTATTTGTTTATCCCCAAGGCGCGTGTAAAGGGGCTTTTCACGGTGGACTGCGC
AmpR promoter
f1 ori

CCCTGTAGCGGCGCATTAAGCGCGGCGGGTGTGGTGGTTACGCGCAGCGTGACCGCTACACTTGCCAGCGCCCTAGCGCC 5440
GGGACATCGCCGCGTAATTCGCGCCGCCCACACCACCAATGCGCGTCGCACTGGCGATGTGAACGGTCGCGGGATCGCGG
f1 ori

CGCTCCTTTCGCTTTCTTCCCTTCCTTTCTCGCCACGTTCGCCGGCTTTCCCCGTCAAGCTCTAAATCGGGGGCTCCCTT 5520
GCGAGGAAAGCGAAAGAAGGGAAGGAAAGAGCGGTGCAAGCGGCCGAAAGGGGCAGTTCGAGATTTAGCCCCCGAGGGAA
f1 ori

DraIII
TAGGGTTCCGATTTAGTGCTTTACGGCACCTCGACCCCAAAAAACTTGATTAGGGTGATGGTTCACGTAGTGGGCCATCG 5600
ATCCCAAGGCTAAATCACGAAATGCCGTGGAGCTGGGGTTTTTTGAACTAATCCCACTACCAAGTGCATCACCCGGTAGC
f1 ori

CCCTGATAGACGGTTTTTCGCCCTTTGACGTTGGAGTCCACGTTCTTTAATAGTGGACTCTTGTTCCAAACTGGAACAAC 5680
GGGACTATCTGCCAAAAAGCGGGAAACTGCAACCTCAGGTGCAAGAAATTATCACCTGAGAACAAGGTTTGACCTTGTTG
f1 ori

PsiI
ACTCAACCCTATCTCGGTCTATTCTTTTGATTTATAAGGGATTTTGCCGATTTCGGCCTATTGGTTAAAAAATGAGCTGA 5760
TGAGTTGGGATAGAGCCAGATAAGAAAACTAAATATTCCCTAAAACGGCTAAAGCCGGATAACCAATTTTTTACTCGACT
f1 ori

TTTAACAAAAATTTAACGCGAATTTT 3′
... 5786
AAATTGTTTTTAAATTGCGCTTAAAA 5′
f1 ori
